# Prophage activity shapes the thermal ecology and evolution of a marine bacterial host

**DOI:** 10.64898/2026.02.19.706829

**Authors:** Catherine A. Hernandez, Jungbin Cha, Noah S. B. Houpt, Jyot D. Antani, Paul E. Turner

## Abstract

Predicting the response of populations to thermal change is challenging due to temperature-dependent interspecific interactions, such as those between bacteria and their integrated viruses (prophages). Temperature is known to modulate prophage activity, raising the question of whether and how prophages alter host responses to temperature. In this work, we explored how prophage infection shapes host thermal ecology and evolution in a novel marine bacterial isolate carrying a spontaneously inducing prophage. We first characterized the effect of temperature on bacteria and phage population dynamics, finding that high temperatures led to bacterial population crashes and reduced yield, with abundant phage production across all temperatures. To then determine the causal impacts of this phage, we infected a conspecific susceptible strain and found that performance in the novel lysogen was reduced across all temperatures and indistinguishable from the naturally infected strain. Further, we found that evolutionary rescue mutants of the naturally infected strain with an expanded upper thermal limit gained high temperature adaptation through a reduction in prophage activity. Overall, we found that intraspecific variation in thermal ecology was explained by prophage infection, and that adaptation to nonpermissive temperature occurred through modifications to prophage dynamics. This work demonstrates a critical link between viral mobile genetic elements and bacterial thermal responses, and emphasizes the importance of ecological interactions in thermal ecology and evolution.

## Introduction

Understanding how organisms respond to changes in temperature is crucial for predicting the impacts of global climate change on biological communities (Dal Bello and Abreu 2024; Huey et al. 2012; Walther et al. 2002; Williams et al. 2008). A useful framework for characterizing organismal responses to temperature is the thermal performance curve (TPC), which describes the relationship between temperature and a metric of organismal performance (Huey and Stevenson 1979). The wide adoption of this framework has generated a rich body of knowledge across a diversity of species and habitats, revealing biogeographic trends such as a latitudinal gradient in the thermal optima of marine phytoplankton (Thomas et al. 2012), and narrower thermal tolerance of ectotherms in the tropics relative to temperate zones (Deutsch et al. 2008; Sunday et al. 2010). However, while useful, it is non-trivial to translate empirical TPC measurements into predictions of fitness across temperatures given the complexities of the biological world (Sinclair et al. 2016). In particular, TPCs are often measured by isolating individuals (or in the case of microbes, strains) and quantifying performance independent of observable interspecific interactions (e.g., in macroorganisms: Kingsolver and Woods 1997; Muñoz et al. 2014; Nagai and Makino 2009; and in microbes: Eng et al. 2023; Ratkowsky et al. 1983; Sieburth 1967). The ecological relevance of this approach is unclear, as interactions between predator and prey (Luhring and DeLong 2016; Miloch et al. 2024), plants and herbivores (O’Connor 2009; Paudel et al. 2020), and interspecific competitors (Tsai et al. 2020; Walberg 2024) have all been shown to impact thermal responses. Even without these easily observable interactions, unseen biotic interactions (such as with endosymbionts or internal parasites) can also alter host responses to temperature, making it challenging to identify the genetic basis of variation in thermal traits (Bakewell et al. 2025; Dunbar et al. 2007; Macnab and Barber 2012). Given the importance of interspecific interactions, characterizing their effects on thermal responses is therefore critical for moving towards a more nuanced understanding of organismal fitness across temperatures.

Microorganisms often live in diverse communities replete with context-dependent interspecific interactions. In particular, bacteriophages (phages; viruses that infect bacteria) are important drivers of bacterial diversity and function across ecosystems (Chevallereau et al. 2022; Koskella et al. 2022), and the effects they have on bacteria depend on environmental factors like temperature (Greenrod et al. 2025). For example, Padfield et al. demonstrated that the addition of a virulent (obligate killer) phage altered the shape of the host population growth rate TPC (Padfield et al. 2019). Beyond virulent phages, bacteria are also frequently infected with integrated viruses called prophages (López-Leal et al. 2022; Touchon et al. 2016). Temperate prophages can either remain dormant within the genome and be inherited vertically in the lysogenic cycle, or become active (induce) and transmit horizontally through the lytic cycle (Lwoff 1953). These replication cycles have drastically different outcomes for individual host cells and whole bacterial populations; in their integrated state, prophages may be neutral or even beneficial for host fitness, while lytic replication leads to production of viral progeny and ultimately cell death (Bondy-Denomy and Davidson 2014; Harrison and Brockhurst 2017). Importantly, prophage replication cycle decision-making is responsive to environmental context. Temperature is one of many known environmental modulators of prophage activity, with high temperature leading to increased lytic replication in many, but not all, studied systems (Bertani and Nice 1954; Kohm et al. 2023; Lieb 1966; Shan et al. 2014; Zeng et al. 2016). This suggests that bacterial thermal responses may depend on both host physiology as well as temperature-dependent prophage activity. However, the effects of prophage infection on host thermal performance and the subsequent consequences for thermal adaptation remain unknown.

In this work, we isolated and characterized a prophage-infected environmental bacterium to determine the consequences of prophage carriage on host thermal ecology and evolution. First, we generated a library of marine bacteria in the genus *Pseudoalteromonas* and screened for spontaneously inducing or inducible prophages. After identifying a focal strain of interest carrying a highly active prophage, we explored how bacteria and phage population dynamics in this strain vary with temperature. Next, to determine the causal effects of this phage on host thermal responses, we integrated the phage into a susceptible conspecific strain, where we could directly measure thermal responses in the presence or absence of the prophage. Finally, we used an evolutionary rescue approach to generate mutants of our original prophage-carrying strain with an expanded upper thermal limit and explored whether improved high temperature growth involved changes to prophage activity. Our work demonstrates the importance of prophage activity in the response of bacterial populations to changes in temperature. More broadly, we add to the growing body of knowledge suggesting that the thermal ecology and evolution of organisms cannot be understood without considering ecological interactions.

## Results

### A marine bacterial isolate carries a spontaneously inducing PM2-like prophage

Through isolating and screening a set of environmental *Pseudoalteromonas* (PSA) strains for phage activity (sampled from coastal waters in the Long Island Sound; see Supp. Fig. 1 for additional details), we identified a strain (CAH24) carrying a spontaneously active prophage that plaques on (i.e., productively infects) a second strain in our library that lacks this prophage (CAH30, Fig. 1a). Using reads from both short- and long-read sequencing, we assembled, annotated, and taxonomically classified closed genomes of both bacterial strains (Supp. Table 1). Each genome is bipartite and contains a 3.69 Mb chromosome and ∼800 kb chromid, and both are closely related to a PSA *shioyasakiensis* genome (GCF_019134595.1; FAST ANI 91.7% for both genomes, alignment fraction 0.796 for CAH24 and 0.83 for CAH30). The genomes of our isolates are highly similar to each other, with 99.93% average identity across aligned regions, although the prophage-carrying strain contains an additional 200kb plasmid-like contig not found in the phage-susceptible strain. This sequence similarity suggests that our two isolates should be considered strains of the same species (Goris et al. 2007; Jain et al. 2018).

**Figure 1.**
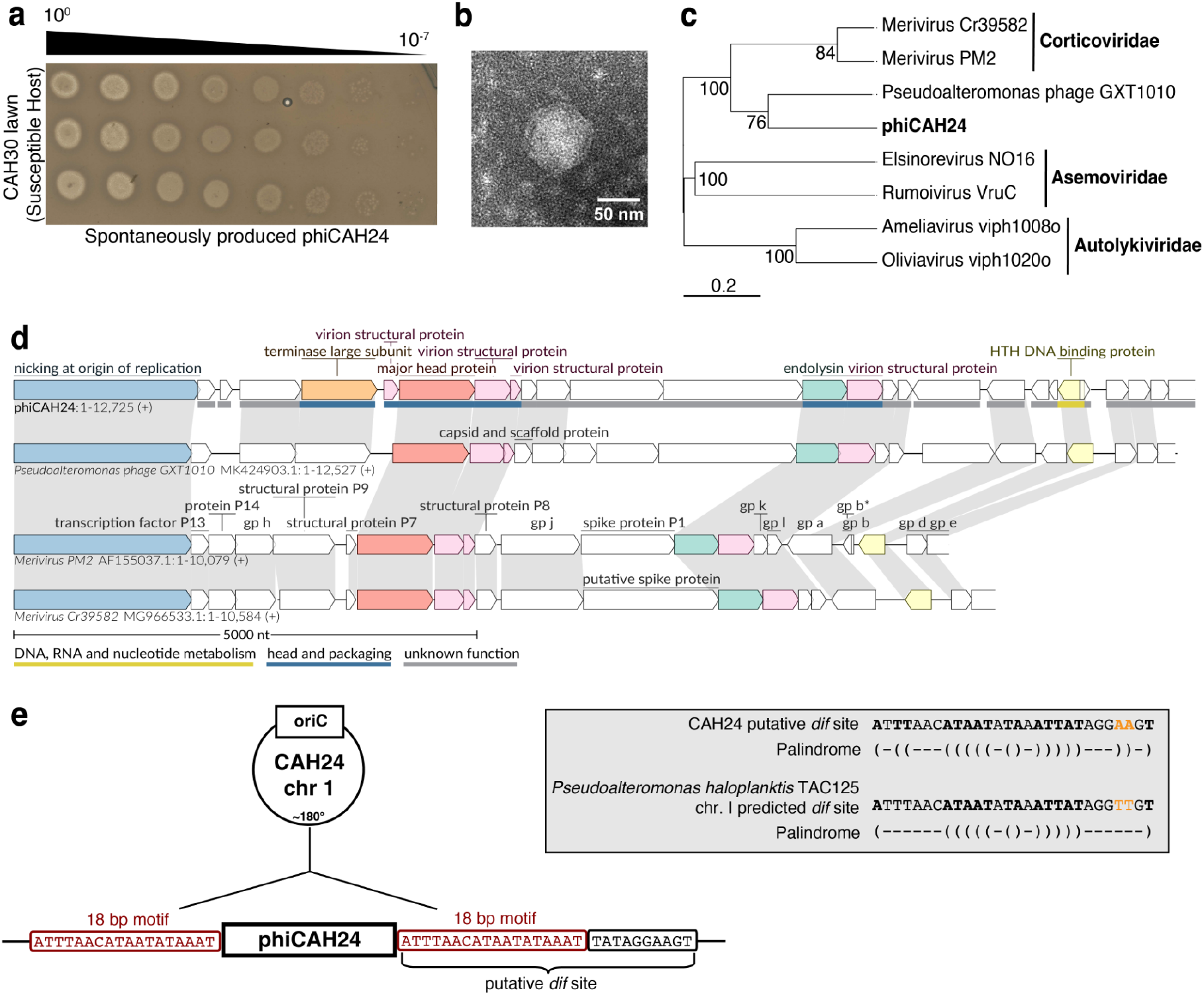
Wild type CAH24 carries a spontaneously inducing, non-tailed PM2-like prophage (phiCAH24). **(a)** Dilution series of spontaneously produced phage from CAH24 spotted on a lawn of CAH30, with the upper black triangle indicating the dilution of the phage sample. **(b)** Representative transmission electron micrograph of spontaneously produced phage, with the scale bar indicating 50 nm. **(c)** Genome-based phylogenetic tree generated by the prokaryotic virus classification tool VICTOR comparing phiCAH24 to other non-tailed phages in the Corticoviridae, Asemoviridae, and Autolykiviridae (based on 2024 ICTV classification). Note that GXT1010 was listed as unclassified by ICTV. Intergenomic distances were calculated using the D6 formula, with branch support values inferred from 100 pseudo-bootstrap replicates. **(d)** Genomic comparison of our phage to existing NCBI PSA Corticoviridae genomes generated using the tool lovis4u. Gene blocks contain a color if there was a functional prediction for that gene in the phiCAH24 annotation from Pharokka, with gray shaded regions connecting homologous protein groups across genomes as defined by MMSeqs2 clustering through lovis4u. **(e)** Evidence supporting the potential for phiCAH24 to be an IMEX (integrative mobile element exploiting Xer recombination), showing its integration position on the replicon relative to oriC, and features of a putative *dif* site directly adjacent to the prophage (TAC125 *dif* prediction from Kono et al. 2011). Parentheses depict complementary base pairs for positions equidistant from the center, reflecting the partial palindromic nature of *dif* sites. Note that one copy of the 18bp motif is present in the circularized form of the phage genome.

Using a combined bioinformatic and morphology-guided approach, we identified the active phage as a non-tailed PM2-like prophage. Two virus prediction tools independently identified two putative prophage regions in the CAH24 genome, one with and the other without predicted tail-associated genes (the “intact” and “questionable” regions in Supp. Fig. 2, respectively). By sequencing the filtrate of an overnight culture and quantifying coverage in our bacterial short-read sequencing, we identified the non-tailed prophage as spontaneously active. All filtrate reads (300x coverage) mapped to the non-tailed region, and short read coverage was approximately 100x greater than in the flanking bacterial regions (Supp. Fig. 2). Transmission electron microscopy supported this molecular result, as we observed non-tailed viral particles of 74 ± 5 nm size in filtrate from a CAH24 overnight culture (mean ± st. dev., n=8 measured particles; Fig. 1b). Top BLAST hits of the predicted genes in this phage identified many matches to the PSA phages PM2 (NC_000867; a model lipid-containing virus, Espejo and Canelo 1968) and Cr39582 (NC_042121), which are both dsDNA phages in the non-tailed Corticoviridae family. We then used a whole genome-based phylogenetic approach to compare our phage to existing NCBI PSA Corticoviridae genomes, as well as two genomes each of non-tailed phages from other families (the Asemoviridae and Autolykiviridae; Fig. 1c, 1d), finding that our phage clustered within a clade containing the currently recognized Corticoviridae phages and phage GXT1010. In summary, we isolated a PSA strain CAH24 (which we also call the wild type, “WT” throughout) carrying a spontaneously active PM2-like prophage (phiCAH24) that productively infects another PSA strain CAH30 (the susceptible host or “SH”) in our library.

### Genomic context of phiCAH24 suggests an IMEX-like strategy

Many prophages encode molecular machinery for integration and excision of their genome from the host chromosome, namely integrases and accessory excisionases. Our annotation of phiCAH24 did not identify any putative integrase or excisionase genes, prompting us to question the possible integration and excision mechanisms for this phage. We found that the genomic context of this prophage shares features with a group of mobile elements called IMEXes (integrative mobile elements exploiting Xer recombination). These elements leverage host recombinases whose typical function is to resolve chromosome dimers by acting on “*dif*” sites in the terminus region of the replicon (Das et al. 2013). We found that phiCAH24 is integrated ∼180° from the predicted chromosomal origin of replication, with an 18bp flanking sequence that is part of a possible *dif* site on one side of the prophage (Fig. 1e). This putative *dif* site is a partial palindrome and differs by 2 bp from a predicted *dif* site in another PSA species (Kono et al. 2011; Fig. 1e). These findings suggest that phiCAH24 may co-opt host enzymes for lysis-lysogeny transitions.

### High temperatures cause CAH24 population crashes and poor performance, but phage activity occurs across temperatures

To explore the relationship between temperature and the performance of CAH24, we first grew replicate populations of the bacterium across temperatures in microplate growth curves. We found that maximum growth rate did not vary with temperature (F_7,2_=7.95, p=0.12), whereas minimum growth rate (indicative of population crashes) and final density were temperature-dependent (F_7,14.27_=50.08, p<0.0001; F_7,7.54_=187.33, p<0.0001, respectively; Supp. Fig. 3). This suggests that temperature did not primarily impact these populations through changes in early growth potential, but instead through driving population crashes and/or reduced final yield. Given this pattern, we chose to use area under the curve, which is an integrated measure that is sensitive to population crashes and final yield, as our population-level measure of “performance”. Plotting these results as a thermal performance curve reveals that CAH24 performance generally declines with increasing temperature, suggesting that this temperature range captures the right side of the TPC for this organism (approximately thermal optimum to the upper thermal limit; Fig. 2b).

**Figure 2.**
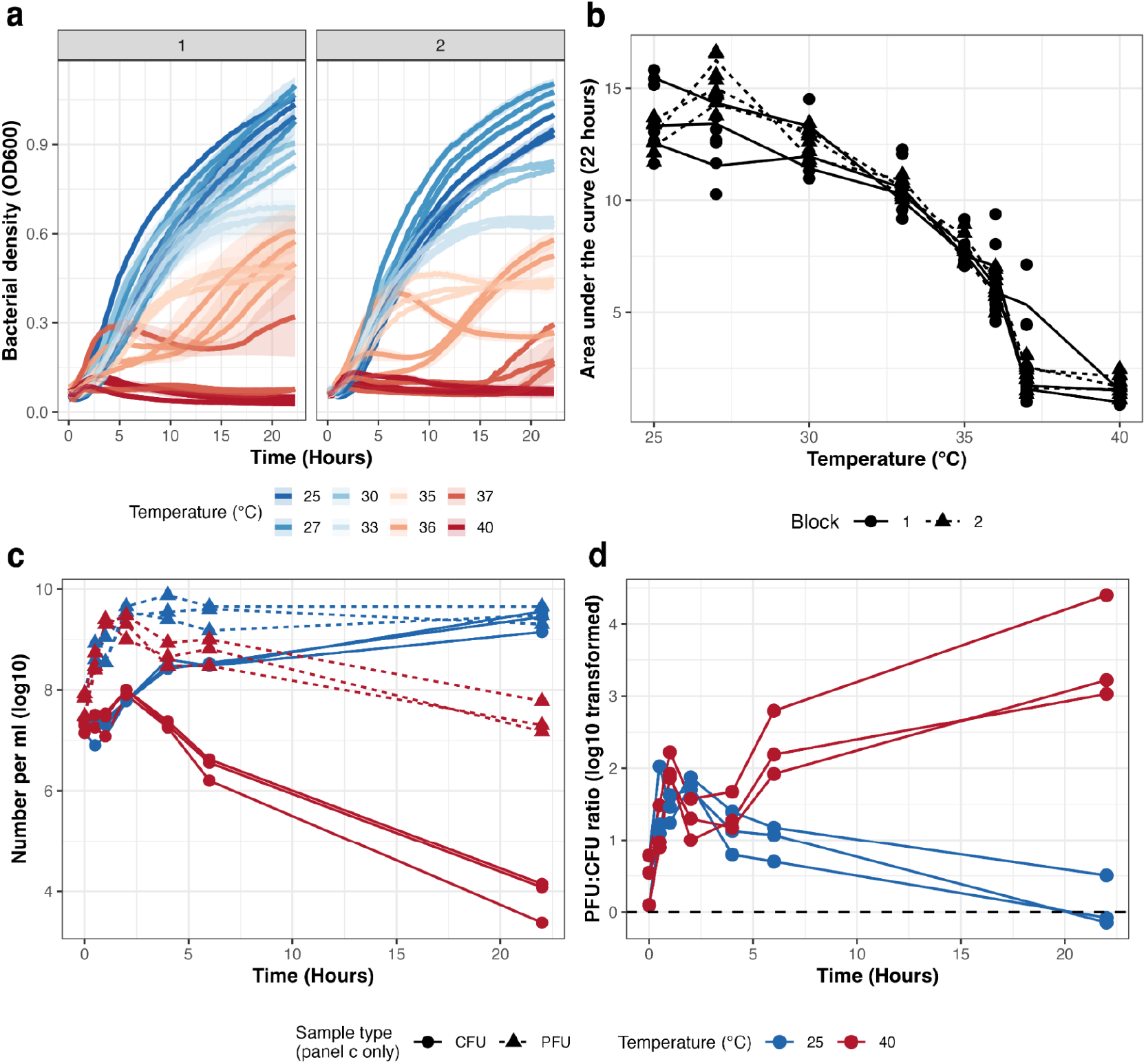
Wild type CAH24 bacterial populations crash at high temperature, while phage is produced at both thermal extremes. **(a)** Bacterial density over time and across a range of temperatures in a microplate reader, with the facets representing two independent blocks of the assay. Bold lines represent the means of each biological replicate (each derived from a unique colony), with the shaded area representing ± SE. **(b)** Thermal performance curve using the area under the curve for each biological replicate across temperature. Points represent the individual values for each well, with lines connecting the means for each biological replicate at each temperature. **(c)** Bacteria and phage population sizes over time at the thermal extremes, as determined by plating for colony forming units (CFUs) and plaque forming units (PFUs). Lines connect points derived from the same initial starting culture, derived from independent colonies of the WT strain. **(d)** The phage:bacteria ratio over time calculated from the data in panel **c**.

To determine the effects of temperature on the combined dynamics of CAH24 and its active prophage, we then grew this strain at our two thermal extremes and measured population sizes across time using a plating-based approach. We found that the bacterial population dynamics qualitatively recapitulated our optical density-based curves, with sustained growth up to a high density at 25°C, and a brief growth phase followed by a decrease in population size after about two hours at 40°C (Fig. 2c). Unexpectedly, we observed high phage titers after the first few hours at both temperatures, suggesting that large amounts of phage were produced across the tested thermal range in this system. We note that cultures also began with some amount of free phage due to carryover from stationary phase cultures. The dynamics at the two temperatures diverged during the later phase of the time series, where the phage titer at 25°C remained about constant, while there was a roughly 100-fold decrease at 40°C. However, the bacterial population decline outpaced that of the phage at high temperature, leading to an increase in the phage:bacteria ratio over time (reaching ≥1000:1), while the ratio decreased and approached 1:1 at the lower temperature (Fig. 2d).

A high PFU:CFU ratio could simply reflect differences in phage decay and non-phage-mediated bacterial death rates at high temperature, or could suggest phage production as the cells are dying. To further probe these possibilities, we asked whether the phage dynamics post-peak (approximately two hours) could be solely explained by decay. To test this hypothesis, we repeated the plating-based time series across three temperatures (25, 38, and 40°C), but for a subset of samples we removed cells by filtration after the estimated peak phage titer to measure decay in the absence of bacteria. From this, we found that decay was temperature-dependent and approximately log-linear at the higher temperatures (temperature*time interaction: F_2,57.03_=102.67, p<0.0001; Supp Fig. 4). Infectious phage were relatively stable at 25°C (-0.008 log_10_ PFU/hr, 95% CI: -.02, 0.005), with intermediate and rapid decay at 38 and 40°C, respectively (38C: -0.06 log_10_ PFU/hr, 95% CI: -0.07, -0.04; 40C: -0.13 log_10_ PFU/hr, 95% CI: -0.15, -0.12). We then used these rates to estimate the 24-hour phage titers expected if phage were simply decaying in the unfiltered cultures during the late phase of the growth curve (between 6 and 24 hours), finding that observed titers were significantly greater than expected at 38 and 40°C (V=19, p=0.047; V=21, p=0.018, respectively), but not at 25°C (V=16, p=0.15). These results are consistent with additional phage production at higher temperatures during the late phase of the growth curve, despite overall declining titers.

### Prophage infection reduces bacterial performance, especially at higher temperatures

To determine the causal effects of this phage on host thermal performance, we infected a closely related strain (the SH, CAH30) with phiCAH24 and compared growth of the uninfected and infected strains across temperature (Supp. Fig. 5). In CAH30, phiCAH24 integrated adjacent to a putative *dif* site homologous to the prophage-adjacent putative *dif* site in CAH24 (Fig. 3a), generating a novel lysogen (a prophage-infected bacterium; which we call the SH-Lysogen or “SH-L”). We next measured spontaneous phage production in the new lysogen and compared its thermal performance to that of its uninfected ancestor. We found that spontaneous phage production by SH-L is similar to CAH24, with over 10^10^ infectious phage per milliliter produced in overnight cultures (WT vs. SH-L contrast p=1; Fig. 3b). In microplate growth curves, the newly infected host had reduced performance across all temperatures relative to its uninfected ancestor (p<0.0001 for all temperatures; Fig. 3c). Strikingly, performance of the two prophage-infected strains (the WT and SH-L) was statistically indistinguishable across the tested thermal range (p>0.3 for all; Fig. 3c), indicating that variation in prophage infection explained the difference in thermal performance of our environmental isolates. While there was an absolute reduction in AUC across temperature for the new lysogen, we also found that the relative reduction in AUC was temperature-dependent, with the relative cost magnified at high temperatures (F_7,25.83_=128.37, p<0.0001; Fig. 3d).

**Figure 3.**
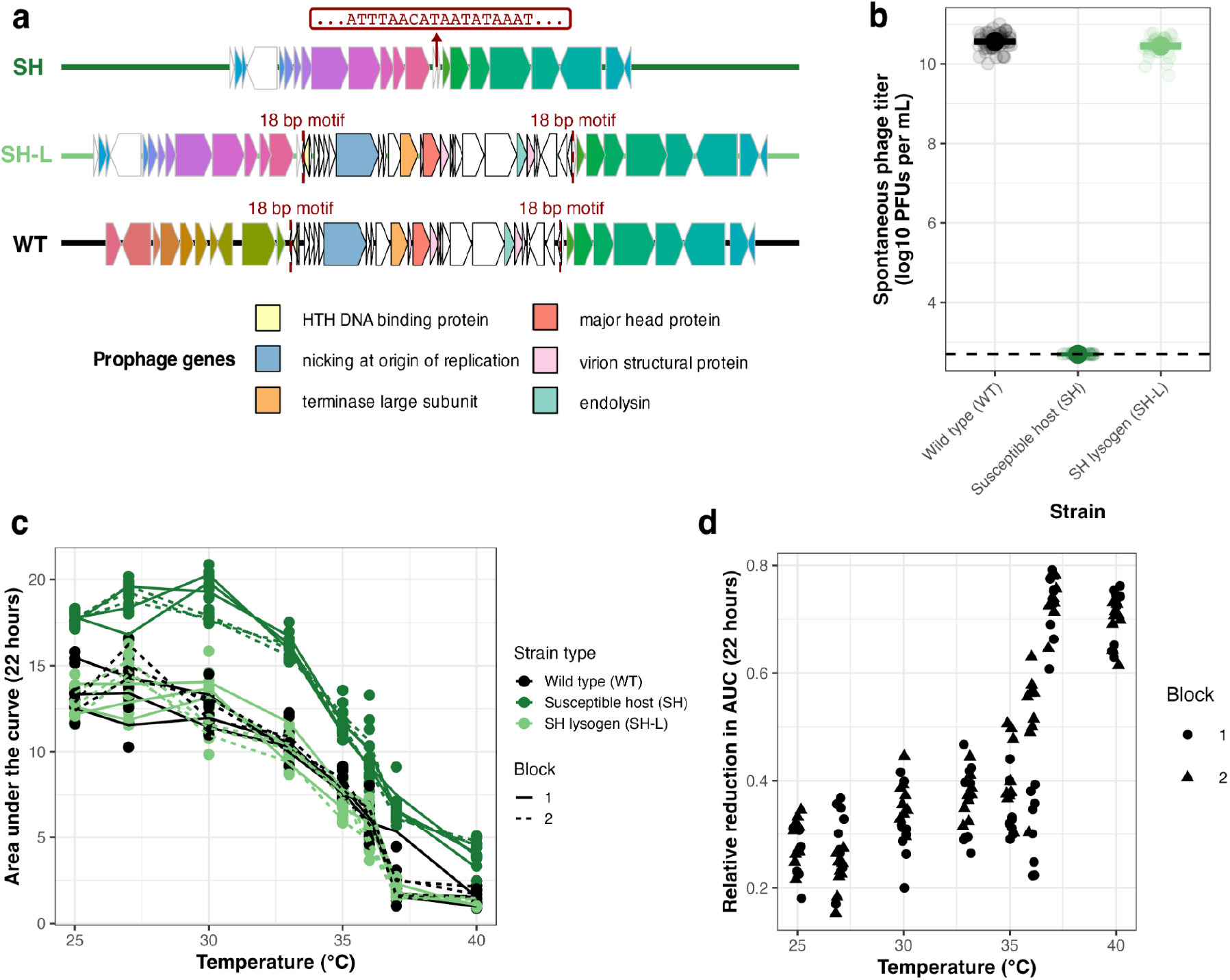
Infection with a spontaneously active prophage reduces performance across temperatures. **(a)** Representations of the SH (susceptible host), SH-L (lysogen in susceptible host background), and WT (wild type) strain genomes in the region of the prophage. Gene blocks with the same color represent genes with the same functional annotation; note that the WT and SH share homology on the right side of the prophage integration site, but not on the left. The red call out box in the SH marks the location and sequence of an 18 bp motif that flanks the phage genome on either side in its integrated state (within SH-L and WT). The legend colors indicate functional predictions for genes found only within the prophage boundary (between the 18 bp motifs). **(b)** Spontaneously produced phage titers (determined by plating on the SH) after 24 hours of growth in marine broth at the original strain isolation temperature (22°C). Each strain was plated across at least six days. Data are summarized as mean ± SE. **(c)** Thermal performance curves of the original prophage-carrying wild type strain, the uninfected susceptible host, and the newly infected lysogen, derived from 22-hour growth curves performed in two independent blocks. For each block, three colonies of each strain were used to found three replicate wells in each growth curve. Points represent the individual values for each well, with lines connecting the means for each colony at each temperature. Note that the WT data originate from the same dataset depicted in Fig. 2. **(d)** Relative reduction in AUC when carrying the prophage (calculated from panel **c** for each temperature as 1 - (AUC_SH-L_ / Avg AUC_SH_)). Higher values represent a greater reduction in AUC due to phage.

### High temperature rescue mutants have reduced phage activity

Finally, we sought to test whether adaptation to high temperature results in altered host-phage interactions in this system. If phage-mediated mortality constrains growth at high temperature, we predicted that mutants selected for growth at high temperature would have reduced phage activity. To test this prediction, we first selected for mutants using an evolutionary rescue approach, where replicate populations of wild type CAH24 were incubated at a temperature just past the estimated upper thermal limit (38°C; Fig. 4a, Supp. Fig. 6). We then monitored the wells over 48 hours for regrowth, and turbid wells were streaked three times to generate pure cultures of putative high temperature mutants (one per turbid well). After an initial screening, three isolates (derived from three initial independent colonies of the WT, and hereafter called the “high temperature mutants” or HTMs) demonstrated robust and improved growth relative to the wild type at the selection temperature, and were chosen for further phenotyping and genetic sequencing (Supp. Fig. 6).

**Figure 4.**
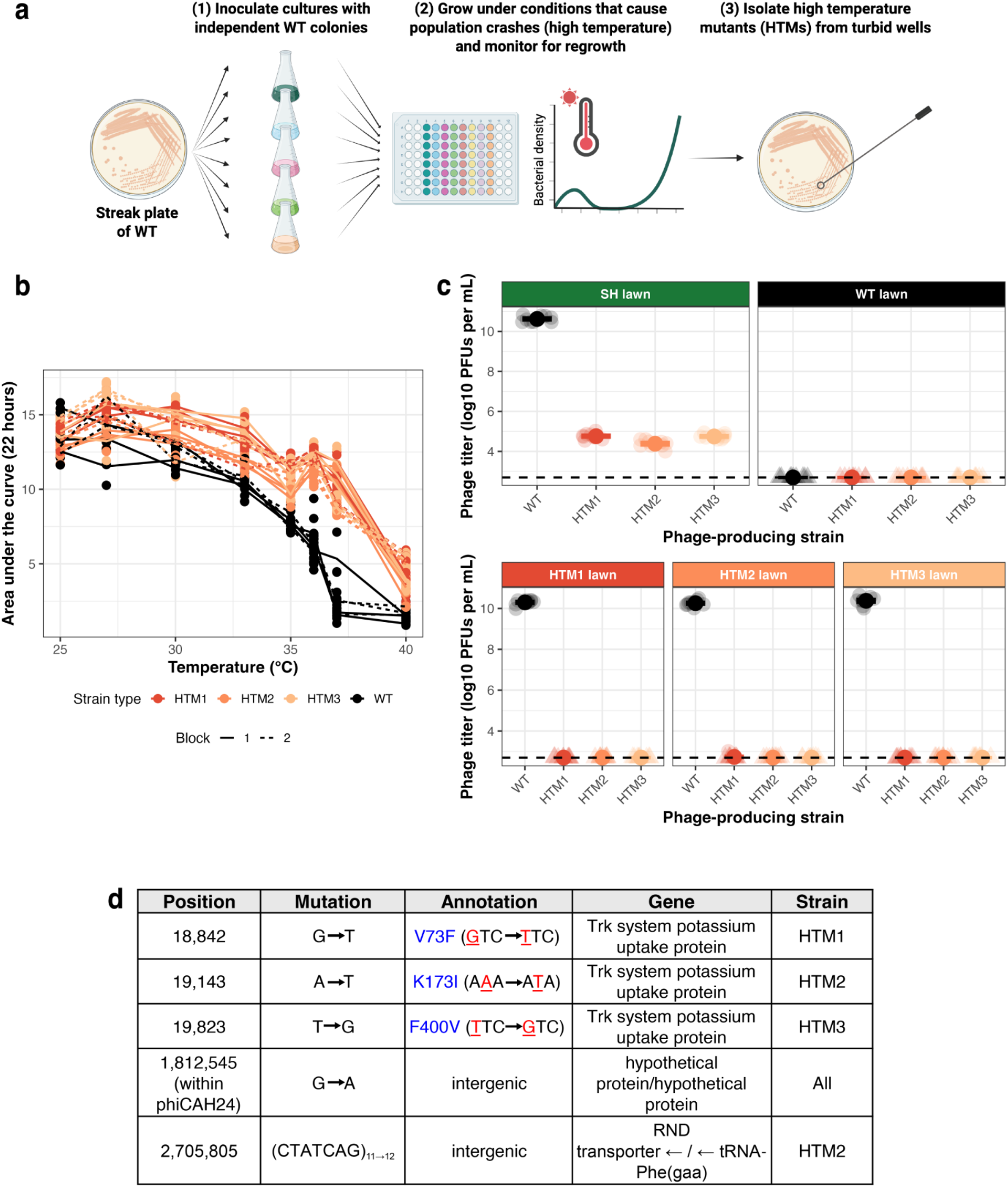
Mutants selected for growth at high temperature have reduced phage production and altered phage susceptibility. **(a)** Schematic representing the methods for generating the high temperature mutants (HTMs). Figure was generated using BioRender (https://BioRender.com/mtgr4sy). **(b)** Thermal performance curves of each HTM relative to their ancestral WT, derived from 22-hour growth curves performed in two independent blocks. For each block, three colonies of each strain were used to found three replicate wells in each growth curve. Points represent the individual values for each well, with lines connecting the means for each colony at each temperature. Note that the WT data originate from the same dataset depicted in Figs. 2 and 3. **(c)** Spontaneously produced phage titers measured on different host strains. Titers of WT and HTM phage were determined on the SH, WT, and each HTM, with data summarized as mean ± SE. For meaningful comparison we show the results of this assay from a single day, but HTM PFUs on the SH were quantified across six days (full dataset shown in Supp. Fig. 7) **(d)** Position and annotations for mutations found in each HTM. Note that all mutations are on the major chromosome, and that the sole shared mutation across strains is within the phiCAH24 region.

We then generated TPCs and assessed phage-related phenotypes of these mutants. We found that all mutants had significantly improved performance from 35-40°C (all p<0.001), but all mutant curves converged with the wild type at low temperature (25°C all p>0.77; Fig. 4b). We then measured spontaneous phage production in overnight cultures and found that all three mutants had significantly reduced phage activity relative to the wild type (WT vs. each HTM plated on the SH, all p<0.0001; full dataset in Supp Fig. 7 with subset shown in Fig. 4c). This result was also supported by reduced sequencing read coverage in the prophage region relative to the wild type (Supp. Fig. 8, compared to Supp. Fig. 2). To determine whether this low phage production was transient or extended to the subsequent growth cycle (given that the temperature curves in Fig. 4b are derived from a second round of growth), we then quantified viable bacteria and phage over time at 25 and 40°C after an initial 24-hour culture. Phage titers generally remained low at both temperatures, often many orders of magnitude lower than the bacterial population size, although there was some variation across replicate wells within a strain (Supp. Fig. 9). Overall, we found that improved high temperature growth was associated with a reduction in prophage activity.

In addition to reduced phage production, the mutants also had altered susceptibility to superinfection (infection of an already infected host) by the WT phage. While the WT phage does not plaque on the WT host, all HTMs were susceptible to the WT phage, although they had reduced susceptibility relative to the SH (all p<0.01; Fig. 4c). HTM phages do not plaque on HTMs or the WT strain. Together, these results suggest that the high temperature-adapted mutants had significantly altered phage-related phenotypes (production and superinfection exclusion patterns).

### An intergenic prophage mutation is associated with adaptation to high temperature

We sequenced our high temperature mutants to determine the genetic basis of our observed phenotypic changes. Whole genome sequencing revealed both shared and unique mutations across the three mutants. They all share the same base pair change in an intergenic region within the prophage (at bacterial genome position 1,812,545), and they each also have a second, unique nonsynonymous substitution in a gene encoding a predicted potassium uptake protein (Fig. 4d). Mutant HTM2 also has a third mutation in an intergenic region. While the presence of multiple mutations in each HTM prevents us from determining the phenotypic effects of each mutation individually without further genetic characterization, we serendipitously observed the emergence of “low inducing” mutant (LIM) colonies from streaks of our frozen wild type strain (Supp. Fig. 10). These colonies produce an intermediate level of phage between the high inducing WT and the HTMs (all p<0.0001, Supp. Fig. 10), and are intermediate in their performance at some high temperatures (LIMs compared to HTMs and WT at 36 and 37°C; all p<0.001; Supp. Fig. 10). Sequencing of six low-inducing and six high-inducing (i.e., wild type) cultures revealed that the LIMs share the same intergenic prophage mutation as the HTMs, with no other predicted consensus mutations in each culture, suggesting that the potassium-associated mutations likely play a role in further reducing phage production and increasing high-temperature performance. While spontaneous plaques within the LIM lawns precluded effective quantification of phage titers on this strain, the prophage mutation in the LIMs qualitatively appears to be sufficient for conferring susceptibility to superinfection by the WT phage (Supp. Fig. 10). These results suggest that a prophage-specific mutation altered superinfection susceptibility, phage activity, and thermal performance, with secondary mutations further modifying the latter two phenotypes.

## Discussion

Thermal responses can be shaped by interactions with other species, but the role of prophage infections in shaping bacterial thermal responses remains unclear. Here, we show that prophage activity underlies both thermal performance and adaptation to high temperature in a species of marine bacteria. By linking prophage activity and thermal performance, our work raises the question of how prophages, and environmentally-sensitive mobile genetic elements more broadly, shape the response of bacterial communities to changing environments.

We found that infection with a spontaneously active prophage significantly altered host thermal performance. Specifically, infection of a marine bacterium with a prophage derived from a conspecific strain caused a downshift in its TPC, including a reduction in the upper thermal limit by at least 3°C. At high temperatures, the greater relative cost of prophage carriage as well as sustained phage production throughout the incubation period suggest that phage activity plays an important role in constraining host population growth. Strikingly, we also found that variation in the TPC between our two strains was completely explained by the prophage, as infecting the naturally uninfected strain led to TPCs that were statistically indistinguishable. This result implies that, in our system, intraspecific variation in thermal ecology across natural isolates was determined by prophage infection status, not variation in host physiology. Across bacteria, thermal traits are emergent properties of complex cellular systems with a polygenic basis (e.g., Roncarati and Scarlato 2017; Siliakus et al. 2017), and show shallow phylogenetic conservatism relative to other traits like pH and salinity preferences (Martiny et al. 2015). Our results suggest that prophages can contribute to this pattern, with horizontal exchange of these elements expected to drive rapid shifts in thermal traits within bacterial communities. Considering the widespread carriage of prophages in bacterial genomes (López-Leal et al. 2022; Touchon et al. 2016), including the Corticoviridae and other non-tailed phage lineages in marine bacteria (Kalatzis et al. 2023; Kauffman et al. 2018; Krupovič and Bamford 2007; Steensen et al. 2024), we expect that prophage-impacted thermal responses are widely relevant in microbial communities.

While our work identified a system with widespread prophage activity and decreases in performance, other prophage-mediated impacts on TPCs are possible depending on mechanism. We hypothesize that beyond this system, the effect of a prophage on host population thermal performance depends on patterns of induction across temperature. For example, prophages with a narrow temperature range between complete lysogeny and widespread lysis may specifically set the upper limit of the TPC rather than cause a downshift across a wider temperature range. These lysis-lysogeny decisions are often regulated by a protein repressor of the lytic cycle, and mutations that render these proteins thermally unstable can lead to high temperature induction (Horiuchi and Inokuchi 1967; Kohm et al. 2023). In such systems, the prophage impact on the TPC will therefore depend on the underlying relationship between temperature and repressor function. However, in systems like ours (a suspected IMEX), the link between temperature and prophage activity may instead arise indirectly through effects on host enzymes involved in chromosomal maintenance during cell division (Xer recombinases; Das et al. 2013). In a non-tailed IMEX of *Escherichia coli*, frequent prophage excision and reintegration were observed even within individual colonies, suggesting widespread activity and instability (Kirchberger and Ochman 2020). Thus, we suspect that cellular division and prophage activity are similarly linked in our system, and this prophage may impose a cost to performance across other environmental factors beyond temperature. Future work should confirm whether this prophage is indeed an IMEX, and explore whether activity occurs under other culture conditions.

Finally, we observed that adaptation to high temperature involved a significant reduction in phage production. This result further supports the model that the upper thermal limit of CAH24 is constrained by prophage-mediated mortality rather than purely host physiology. While these mutants had significantly reduced phage production and improved performance at high temperature, performance was not universally improved across the tested thermal range. Given the extreme reduction in phage production and, presumably, phage-driven mortality across temperatures, this suggests these mutants may suffer from costs to other cellular processes at lower temperatures. Sequencing of the high temperature rescue mutants revealed two shared types of mutations. First, they shared the same base pair change in an intergenic region of the prophage, which in close relative phage PM2 is known to be an important regulatory region with several promoters (Männistö et al. 2003). Evidence from our LIMs suggests that this mutation alone substantially reduced phage production and moderately improved high temperature performance. Second, all HTMs also had nonsynonymous substitutions in a bacterial gene associated with the Trk potassium uptake system. In *Streptococcus pneumoniae*, a mutation in a Trk system gene improved heat shock survival through effects to the intracellular signaling molecule c-di-AMP (Zarrella et al. 2018). While the mechanisms by which these mutations impacted cellular physiology and prophage regulation in our strains remain unknown, their parallel occurrence in the HTMs implies a potential link between thermal adaptation, prophage activity, and ion homeostasis. More broadly, the extent to which evolutionary rescue at extreme temperatures is underlain by changes in host physiology vs. changes in interactions with other species remains an exciting and open area of research.

In the face of global climate change, it is imperative that we uncover the factors shaping the response of populations to increasing temperatures. Here, we demonstrate that prophage infection can significantly reduce thermal performance and the critical thermal maximum of their bacterial hosts. Further, we show that mutations conferring improved performance at high temperature are associated with reduced prophage activity. Given the vast diversity of phages, one could imagine a range of other possible connections between prophages and temperature responses, perhaps with increases to thermal performance or specific impacts on lower thermal limits. By exploring the temperature-prophage connection in our novel environmental strains, these results further justify the importance of incorporating interspecific interactions in thermal ecology and evolution.

## Methods

### Environmental strain isolation and culturing conditions

We sampled tide pools and the nearshore waters of the Long Island Sound at Chaffinch Island Park, Guilford, Connecticut, USA (41.264°N, -72.675°W) at approximately peak low tide on September 9, 2022 and February 28, 2023 to generate a library of marine bacterial strains. For each sample, we collected 40 mL of upper surface water in a sterile 50 mL syringe and filtered through a 0.22 µm MCE membrane (GSWP02500) in an autoclaved Millipore Swinnex Filter Holder (25mm, SX0002500). Time, temperature, and pH were recorded for each sample, with temperature and pH measured using an EcoSense EC300A Conductivity Meter (YSI Inc., Yellow Springs, OH). Filters were stored in sterile petri dishes on ice until brought to the lab for processing (max time on ice about two hours). In the lab, each filter was cut in four pieces using a flame sterilized razor, and ¼ of each filter was placed on a marine agar (Difco™ Marine Agar 2216, BD Diagnostics) plate for incubation at 22°C. After two days, we streaked three colonies (with distinct morphologies when possible) from each sample onto new marine agar plates and incubated again at 22°C. This process was repeated at least three times to generate pure cultures, after which a single colony of each strain was grown overnight in 5 mL marine broth (Difco™ Marine Broth 2216, BD Diagnostics) in test tubes at 22°C with shaking at 200 rpm. Long term stocks of each strain were stored as a 1:1 ratio of overnight culture with 50% glycerol at -80°C.

### Strain identification

To identify each strain at roughly the genus level, we amplified partial 16S rRNA sequences (V3-V4 region, 331F-797R, Nadkarni et al. 2002; Supp. Table 2) using colony PCR. 50 μL reactions were prepared with: 10 μL 5X Green GoTaq Flexi Buffer (Promega, M891A), 4 μL 25 mM MgCl2, 1 μL 10 mM dNTPs, 0.5 μL 10 μM each primer, 0.25 ul GoTaq Flexi DNA Polymerase (Promega, M829A), 33.75 μL molecular grade water, and a small amount of each colony as template. Reaction cycling conditions were as follows: 95°C 5 minutes, 35 cycles of [95°C 30s, 50°C 30s, 72°C 1.5 min], 72°C 15 min, 12°C hold. Amplification was confirmed by running 25 μL of each PCR reaction on a 1% agarose gel in 1x TAE with 15 μL of 1 mg/mL EtBr at 120V for about 30 minutes. Visible bands at the appropriate size were cut and amplicons extracted using the GeneJET Gel Extraction Kit (Thermo Scientific, K0691) following all steps as specified by the manufacturer, except that the samples were eluted in 50 μL molecular grade water. Purified products were submitted for Sanger sequencing at the Keck DNA Sequencing Core (Yale University). After trimming the low quality ends, we compared each resulting sequence to the 16S rRNA database hosted by NCBI using blastn. Strains identified as part of the genus *Pseudoalteromonas* at the time we began prophage hunting (11x September, 17x February) were included in the following experiments.

### Active prophage discovery and phage plating methods

To identify *Pseudoalteromonas* isolates with inducible and/or spontaneously inducing prophages, we incubated all strains in triplicate in the absence and presence of the general chemical inducer mitomycin C (0.5 μg/mL) in 96 well plates at 22°C and shaken at 200 rpm. Note that this method is only expected to capture a subset of all possible inducible prophages (e.g., Dahlman et al. 2025). After 24 hours of incubation, cultures were filtered (0.22 μm) to remove bacterial cells, and diluted 1:100 in marine broth to reduce the amount of mitomycin C carryover. We then spotted 1 ul of each diluted filtrate onto double layer plates containing each of the *Pseudoalteromonas* strains (1 well plate CellTreat 229501, marine agar base, top layer 10 mL marine agar 0.3% with 100 μL overnight culture) and incubated the plates at 22°C. We recorded whether there was any visible clearance in spots as preliminary evidence of phage infection of that host. One filtrate had visible clearance on a single host strain (Supp. Fig. 1), and phage production was subsequently confirmed by plating a dilution series on a double layer plate containing the putative host and checking for individual plaques. For all phage titering in this study, double layer plates contained a base layer of marine agar, and top layers contained a mixture of soft marine agar (0.3% agar) with 1% bacterial overnight culture. Plates were then incubated at 22°C overnight. From these assays, we identified a *Pseudoalteromonas* strain (CAH24; site D, September) carrying a spontaneously inducing phage (phiCAH24) that infects another strain in this library (CAH30; site OCE2, September; Supp. Fig. 1).

### Transmission electron microscopy

We used transmission electron microscopy to visualize phiCAH24. Carbon-coated grids (Ted Pella Inc Carbon Type-B, 400 mesh, Copper, catalog #01814-F) were cleaned with a PELCO easiGlow™ Glow Discharge Cleaning System. 4 μL drops of fresh, spontaneously produced phage stock (∼10^10^ PFU/mL, from a 0.22 μm filtered 22°C overnight culture of WT CAH24) were placed on each grid, and the grids were negatively stained with 2% uranyl acetate. Transmission Electron Microscopy was performed using a 120 kV Talos L120C instrument at the Yale CryoEM Resource facility. Diameters of viral particles were measured in Fiji (ImageJ) using the straight line selection tool and the Analyze > Measure function. The diameter is reported as mean ± standard deviation from 16 total measurements of 8 viral particles.

### Lysogen creation

To test for a causal impact of the phage on host thermal responses, we integrated the phage into the susceptible host (CAH30) genetic background. Three freshly prepared high titer phage filtrates were spotted on a square (100 mm x 100 mm) marine agar plate with a top layer containing 7 mL marine agar (0.3%) and 70 μL overnight culture of the susceptible host. The plate was incubated at 22°C for 24 hours, after which a “mesa” of regrown bacteria was observed in the center of each high titer phage spot. We then streaked for individual colonies from each mesa and screened 12 colonies from each streak (n=36) for phage production by spotting overnight cultures on a top layer containing the original susceptible host (as well as spotting the original wild type strain and susceptible host as controls). One colony had a ring of clearance surrounding its spot and was presumed positive for phage integration and spontaneous production. This colony was restreaked twice more and frozen at -80°C in 25% glycerol for long term storage.

After sequencing and generating a draft genome for this novel lysogen (see details in sequencing section below), we designed primers to confirm integration by amplifying the new host-phage junctions on either side of the prophage (Supp. Table 2). For each primer pair, one primer anneals to a sequence within the prophage genome, and one primer anneals to a sequence on the host genome. The “right” side primers were expected to amplify both the novel lysogen and WT CAH24 due to homology on that side of the integration site, but the “left” primers were expected to amplify only the novel lysogen. We screened six colonies of the novel lysogen for integration using these primers, with a single colony of the WT and SH included for comparison. 50 μL reactions were prepared with: 10 μL 5X Green GoTaq Flexi Buffer (Promega, M891A), 4 μL 25 mM MgCl2, 1 μL 10 mM dNTPs, 0.5 μL 10 μM each primer, 0.25 μL GoTaq Flexi DNA Polymerase (Promega, M829A), 33.75 μL molecular grade water, and a small amount of each colony as template. Reaction cycling conditions were as follows: 95°C 5 minutes, 34 cycles of [95°C 30s, 54°C 30s, 72°C 1 min], 72°C 5 min, 12°C hold. Results were visualized by running 10 μL of each PCR reaction on a 1% agarose gel in 1x TAE with 15 μL of 1 mg/mL EtBr at 120V for about 25 minutes.

### High temperature mutant selection

We performed a high temperature evolutionary rescue experiment to select for mutants able to grow past the wild type CAH24 upper thermal limit. Seven colonies were first grown 24 hours in marine broth at 22°C, then diluted 1:200 in marine broth and each used to begin eight replicate wells in a 96 well plate. This plate was incubated in a BioTek Epoch 2 plate reader at 38°C for 48 hours, with constant shaking and OD600 measurements every 15 minutes. After 48 hours, turbid wells were streaked for single colonies on marine agar plates incubated at 22°C. One colony per turbid well was selected for restreaking and long term storage in glycerol at - 80°C. To determine whether the re-streaked strains were capable of high temperature growth, we again grew them at the selection temperature (38°C) and compared their growth dynamics to their ancestral wild type CAH24 and the susceptible host CAH30. From these experiments, we had five turbid wells after the initial selection plate (3A, 3H, 6A, 7H, 8H; Supp. Fig. 6), with four strains able to grow on subsequent streak plates (well 7H did not produce colonies), and three strains with robust growth at 38°C that were selected for subsequent sequencing and phenotyping (wells 3A, 6A, and 8H, which became the high temperature mutants or “HTMs”; Supp. Fig. 6).

### Sequencing

We used both short- and long-read sequencing to assemble genomes (WT CAH24, SH CAH30, SH-L), and used short-read sequencing to identify variants (HTMs, LIMs). For all strains, DNA for short-read sequencing was extracted from overnight cultures of individual colonies using the Qiagen DNeasy Blood and Tissue Kit (cat no. 69504) following manufacturer instructions for Gram-negative bacteria. Illumina sequencing was performed by SeqCoast Genomics (New Hampshire, USA) using the Illumina NextSeq2000 platform producing 2x150 bp paired reads, with read demultiplexing and trimming performed using DRAGEN v3.10.12 (WT, SH, HTMs) or v4.2.7 (SH-L, LIMs), depending on when the samples were sequenced. To assemble a complete genome of the WT and SH, we shipped streak plates of each strain to SeqCoast Genomics for long-read sequencing along with their recommended in-house DNA extraction protocol. Cells were scraped from a high density area of the plate, lysed using MagMAX Microbiome bead beating tubes, and DNA extracted using the Qiagen DNeasy 96 PowerSoil Pro QIAcube HT Kit protocol. Oxford Nanopore long-read sequencing was performed using R10.4.1 flow cells (WT: on the GridION platform, SH: on the PromethION 2 Solo platform) with ‘super-accurate’ basecalling and barcode trimming enabled. To identify the prophage insertion site in the lysogen, in addition to short read sequencing, we extracted bacterial overnight culture DNA using the Blood and Tissue Kit and long-read sequencing was performed by Plasmidsaurus using an R10.4.1 flow cell and super-accurate basecalling. Mutations in the LIMs were identified by sequencing six overnight cultures each of high inducing WT colonies and low inducing LIM colonies.

As part of our efforts to identify the spontaneously active prophage, we sequenced filtrate from an overnight culture of WT CAH24. We first removed bacteria from 7.5 mL of overnight culture via centrifugation and filtered the supernatant through a 0.22 μm filter. We then precipitated phage particles by adding a PEG and NaCl solution to the filtrate (final concentration 4% PEG 8000, 0.5 M NaCl) and incubating overnight at 4°C. After overnight incubation, the sample was centrifuged at 10,000×g for 30 minutes, and the pellet was resuspended in phage resuspension buffer (1 M NaCl, 10 mM Tris•HCl pH 7.5, 0.1 mM EDTA). We then removed free nucleic acids by treating the sample with DNase I (1 μL, 2000 U/mL) and RNase A (1 μL, 20 mg/mL) and incubating for 30 minutes at 37°C. After adding 10 μL of EDTA (0.5 M, pH 8.0), we then performed a genomic extraction using an equal volume (to the sample) of phenol:chloroform:isoamyl alcohol (25:24:1). The mixture was centrifuged at 16,000×g for 5 minutes, and the resulting aqueous phase was transferred to a new tube. Next, we ethanol-precipitated the DNA by adding 1000 μL of pre-chilled 100% ethanol and 50 μL 3M sodium acetate, and stored the sample at -20°C overnight. The sample was then centrifuged at 14,000×g for 15 minutes, and the pellet resuspended in 500 μL pre-chilled 70% ethanol. After another centrifugation at 14,000xg for 5 minutes and allowing the pellet to air dry, the pellet was resuspended in 50 μL water and incubated at 37°C for 30 minutes. Finally, the DNA was further cleaned by using the Zymo DNA Clean and Concentrator Kit following manufacturer steps, with a final elution in water. This sample was submitted to Plasmidsaurus for long-read sequencing, which used R10.4.1 flow cells and Dorado for basecalling. All sequencing data (raw reads and annotated genomes) will be available under NCBI BioProject PRJNA1417279 upon publication.

### Sequencing analyses

Using our sequencing data, we assembled, annotated, and classified the bacterial strains in this work. For Illumina short reads, trimming was performed by the sequencing company and we used all provided reads that passed their quality control, with positional mean quality near or above 30 at all positions. In bacterial assemblies, Nanopore long reads were primarily used for bridging assembly gaps (using a “short read first” hybrid assembly approach), and all reads passing the basecaller quality filter were included. Hybrid assemblies of the WT (CAH24), SH (CAH30), and SH-L (phiCAH24 integrated in SH background) were created with Unicycler (v0.5.0), producing completed circular contigs for the WT and SH. Mean depth, base quality, and map quality for the WT and SH were determined by mapping the short reads back to each assembled genome using Bowtie 2 (v2.5.4), and calculating assembly statistics using samtools (v1.23, Supp. Table 1). The SH-L assembly was incomplete but contained the prophage on a larger contig with bacterial genes, and thus was sufficient for identification of the insertion site. For downstream analyses, genomes of the WT and SH were reoriented to start at dnaA using circlator (v1.5.5), and annotated using Bakta (v1.11.0). We ran GTDB-Tk (v2.4.1) using reference data version r226 to classify the WT and SH, which identified a top hit that has since been suppressed on NCBI for being too small (GCA_013391845.1), so we report the next best ANI match for both strains. In addition, we ran dnadiff within the MUMmer tool (v3.23) to compare the genomes to each other. To identify mutations in the HTMs and LIMs relative to our annotated WT genome, we ran breseq (v0.38.1; Deatherage and Barrick 2014) in consensus mode and reported all predicted, non-marginal mutations.

We then used various tools to identify predicted and active prophage(s) in the WT CAH24 strain. To predict prophages in the WT strain, the annotated genome was imported into Proksee where we ran PHASTEST (v1.0.1) and Virsorter (v1.1.1), which both identified two predicted prophages. We then used read coverage to identify the spontaneously active prophage in our short read sequencing of a WT overnight culture, aligning the reads to our reference using Bowtie 2 (v2.5.4) and quantifying coverage using samtools (v1.21) in the regions of both predicted prophages. These tools were also used to quantify read coverage in short read sequencing of the HTMs and LIMs. In addition, we long-read sequenced a spontaneously produced phage lysate using Plasmidsaurus (300x coverage), which independently assembled a single phage genome that was a 100% match to one of the predicted prophage regions within our CAH24 assembly. We then ran Pharokka (v1.7.5, using Phanotate v1.5.1) to generate an annotation of this phage using a phage-specific tool. The annotations of this prophage region identified many BLAST hits to Corticoviridae phages, so to further classify this phage we used the web service tool VICTOR (https://victor.dsmz.de), which generates a whole genome-based phylogeny and classifies prokaryotic viruses. In addition to our phage genome, we included all three currently available PSA Corticoviridae genomes in NCBI GenBank (current as of Sep 25, 2025; although note that GXT1010 was listed as unclassified in the ICTV 2024 report), as well as two Autolykiviridae and two Asemoviridae genomes (Kalatzis et al. 2023; Kauffman et al. 2018). Note that the VICTOR classification results did not follow the accepted ICTV classifications for these genomes, and we present the ICTV classification in our results and figures. Finally, we used lovis4u (v0.1.4.1) to visualize synteny and homologous protein groups in our phage relative to the NCBI PSA Corticoviridae phage genomes.

Given the lack of clear integration and excision-specific genes in our phage annotation, we next probed the possibility of this phage relying on host recombination machinery. In particular, some prophages exploit bacterial Xer recombinases (and are classified as IMEXes: integrative mobile elements exploiting Xer recombination). These elements often integrate at “*dif*” sites located approximately 180° from the chromosomal origin of replication (oriC). We compared the Bakta oriC prediction in the annotation of our WT strain to the oriC predicted by DoriC (12.0; Dong et al. 2023) in the PSA RefSeq genome that is closest to this strain (GCF_019134595.1), finding that the DoriC sequence overlaps with the region predicted by Bakta (Bakta: 3,676,243-3,676,646; DoriC: 3,676,240-3,676,668), and they share 95.35% similarity. Next, we searched for a putative *dif* site in the major chromosome by comparing a predicted *dif* site in a congeneric species (*Pseudoalteromonas haloplanktis* TAC125, NC_007481; Kono et al., 2011). The best match to this sequence is directly adjacent to the prophage boundary (positions 1,824,787-1,824,814), differing by 2 bp from the *P. haloplanktis* sequence in a way that increases palindromicity. Taking the midpoint of this region (position 1,824,800), the approximate midpoint of the predicted oriC region (position 3,676,454), and the length of the chromosome (3,686,693 bp) we calculated the distance between the *dif* site and oriC in degrees (180.81°).

### Spontaneous phage production and efficiency of plating assays

Spontaneous phage production was quantified for each strain by inoculating individual colonies into 1.5 mL marine broth in a 24 well plate, shaken at 200 rpm and incubated at the isolation temperature (22°C). After 24 hours of growth, cells were pelleted and the supernatant filtered through a 0.22 μm filter. Samples were diluted in a 1:10 series using marine broth, and then spotted onto marine agar plates with top agar lawns containing 1% overnight culture of the susceptible host CAH30. Plates were incubated at 22°C overnight, and phage titer was determined by counting individual plaques. Phage production was measured for each strain across many independent colonies and days, with replication details in figure legends.

To quantify efficiency of plating (EOP), spontaneously produced phage filtrates were plated on lawns containing hosts of interest, with the susceptible host serving as the point of reference on each assay day. Fresh filtrates were prepared across many replicate days, with consistent qualitative results. For ease of comparison, we plot the results of the most comprehensive dataset we have from a single day in Fig. 4c. On this day, three colonies of each phage-producing strain (WT, LIM, SH-L, and HTMs) were used to generate filtrates that were plated on lawns of all strains, including the susceptible host. Given variable limits of detection due to large differences in titers on the SH, we plot the raw phage titers of each strain on each host in the main text. Images of all plates were scanned using an Epson Perfection V850 Pro scanner. EOPs on LIM lawns were not possible to count accurately due to spontaneous plaques derived from the overnight cultures, but we were able to qualitatively call resistance or susceptibility, with representative images presented in the supplement (Supp. Fig. 10).

### Temperature growth curves (optical density and culture-based)

We quantified growth over time using optical density as a proxy for bacterial population size in a temperature-controlled plate reader. First, we grew cultures of three random colonies from streak plates of each strain to stationary phase (marine broth, 22°C, shaken 200 rpm). After 24 hours of growth, cells were harvested by pelleting 1 mL of each culture at 4000 rcf for 10 minutes, and washed by removing the supernatant and resuspending in 1 mL of fresh marine broth. This process was repeated twice to remove spent media, after which the cells were diluted in marine broth to a starting OD600 of 0.05. We prepared three replicate wells per colony (200 μL) in a flat bottom 96 well plate to measure growth in a plate reader-based assay. All strains (WT, LIM, SH, SH-L and HTMs) were represented on each plate, with well position randomized each day, and outer columns avoided to reduce edge effects. Growth curves were performed using two BioTek Epoch 2 plate readers (Agilent Technologies, California, USA) with the following settings: a single controlled temperature, constant double orbital shaking, and OD600 reads every 15 minutes for 22 hours. We selected eight temperatures spanning the lowest possible in our setup (25°C; approximately the temperature of the tide pool from which we isolated this strain, Supp. Fig. 1) to a temperature with minimal growth of the WT strain, with increased resolution around the upper thermal limit (list of all: 25, 27, 30, 33, 35, 36, 37, 40°C). Growth across all temperatures was measured in two independent blocks, with cultures derived from the same three colonies used to inoculate each growth curve plate within each block (for a total of six unique colonies representing each strain in the full dataset). We assessed the effects of temperature on these curves by calculating observed growth rates (using the slopes of ln(OD600) with a 5 time point rolling window linear regression; the “easy linear” method, Ghenu et al. 2024), and then extracted the maximum and minimum values (after reaching a threshold OD of 0.07, to identify crashes), as well as final density for each temperature. Given the atypical growth dynamics of many strains in these conditions, we then quantified “performance” as area under the curve (using the auc function from gcplyr; Blazanin 2024) in each well. Note that our general approach (growth at a benign temperature, dilution, and immediate incubation at assay temperature) is reflective of acute responses to a rapid temperature shift.

For a subset of strains, we also quantified bacteria and phage titers over time at different temperatures using culture-based methods. Cultures were prepared in the same way as the plate reader growth curves, with stationary phase cells washed twice and resuspended to an OD600 of 0.05. The cultures were then incubated in a 96 well plate with constant 200 rpm shaking in a New Brunswick I26 Incubator Shaker set to a defined temperature. Wells were destructively sampled at each time point for culturable bacteria and phage population sizes. To determine bacterial population sizes, we generated a 1:10 dilution series of each sample at each time point, which were spotted on marine agar plates incubated at 22°C. To quantify phage titers, we first pelleted bacteria (centrifuged at 3220xg for 10 minutes), and then filtered the supernatant through a 96 well 0.22 μm filter plate (Millipore MSGVS2210, centrifuged at 3220xg for two minutes), after which we plated a 1:10 dilution series of filtrate on top layers containing 10% (by volume) overnight culture of susceptible host (plates incubated at 22°C). For a supplementary experiment, we also filtered bacterial cells out of a subset of the incubating wells after two hours and incubated the cell-free phage cultures to determine decay over time under our experimental conditions.

### Statistics and figures

Statistical analyses and figure generation were primarily done in R (v4.5.0) and arranged in Adobe Illustrator, with other programs specified where relevant in the manuscript. Broadly, our analysis approach consisted of generating linear mixed models including fixed effects of interest and blocking factors as random effects. For significant fixed effects, we then performed post-hoc analyses by comparing estimated marginal means between groups of interest, with p-values corrected for multiple comparisons using the Holm method (when comparisons were between planned groups) or Tukey method (when testing all pairwise comparisons).

Thermal performance curves were analyzed as linear mixed models (R package lme4, v1.1-37) testing for a strain by temperature interaction on area under the curve, with random effects for each blocking variable (Block/date/plate, plate reader, colony). We chose not to fit formal thermal performance curve models to our data, as we are only capturing a subset of the curve with our achievable temperature range. The full dataset containing all strains was used to generate the initial overall linear model. Post-hoc comparisons were performed of planned comparisons in two groups using estimated marginal means (emmeans v.2.0.0), the first comparing the WT, SH, and SH-L strains to each other across all temperatures, and the second comparing the WT, LIM, and HTMs to each other. Differences in observed maximum and minimum per-capita growth rates and final density in the wild type strain curves were compared using the same random effects structure, with temperature as the single fixed effect and no post-hoc comparisons. To test for a temperature effect on the relative reduction in AUC due to prophage carriage, we first calculated this value as 1 - (AUC_SH-L_ / Avg AUC_SH_), then built a model including temperature (fixed) with Block/date and colony (random). All pairs of temperatures were then compared using estimated marginal means.

To analyze the results of the wild type plating-based time series with unfiltered and filtered samples, we used phage decay rates from the filtered data to determine whether the patterns in the unfiltered data could be explained by those rates alone. We first tested for a temperature by time interaction with colony as a random effect in the phage decay data. We then used this to determine a single decay slope of log_10_ PFU per mL for each temperature, which was used to calculate the expected PFUs in each unfiltered culture if phages were simply decaying in the late phase of the growth curve (between 6 and 24 hours). We then tested whether the difference in observed and expected log_10_ PFUs per mL was significantly greater than 0 using a one-sided Wilcoxon test for each temperature.

We next analyzed phage plating results following our mixed model approach. Samples producing no detectable phage (below the limit of detection) were excluded from both models. Spontaneous PFU production (log_10_ transformed) of each strain was analyzed as a mixed model testing for a fixed effect of strain with a random effect accounting for day-to-day variation, with estimated marginal means of all strains compared post-hoc. For our superinfection exclusion/efficiency of plating data (where WT and mutant phages were plated on WT, SH, and mutant lawns), we present and statistically analyze a comprehensive dataset of raw titers collected on a single day for meaningful visual comparison. This assay included an all x all cross, where for each strain we had three filtrate-generating colonies plated on lawns derived from three colony overnights, for a total n=9 of each phage×host combination. We tested for a phage×host interaction on log_10_ PFUs per mL, with a random effect for phage-producing colony ID and colony ID in lawn. We then tested whether WT phage titers differed across host strains using their estimated marginal means. We note that many combinations of interest were plated across additional days with consistent qualitative results, with all data and analysis code for this manuscript available upon publication.

## Acknowledgements

We are grateful to Mary Beth Decker and Jennifer Velasquez for help in the field, and all members of the Turner lab (particularly Helen Stone, Albert Vill, and Dallas Mould) for feedback on this work. Funding support to CAH was provided by the Yale Institute for Biospheric Studies Gaylord Donnelley Postdoctoral Environmental Fellowship, and a National Science Foundation Postdoctoral Research Fellowship in Biology (Award No. 2109819). JC was supported by a Yale College First-Year Summer Research Fellowship in the Sciences & Engineering through the Yale Science and Quantitative Reasoning Education department. NSBH was supported by a National Sciences and Engineering Research Council of Canada Postgraduate Scholarship (CGSD3 - 559651 - 2021).

