## Supplementary figures and tables for "Prophage activity shapes the thermal ecology and evolution of a marine bacterial host"

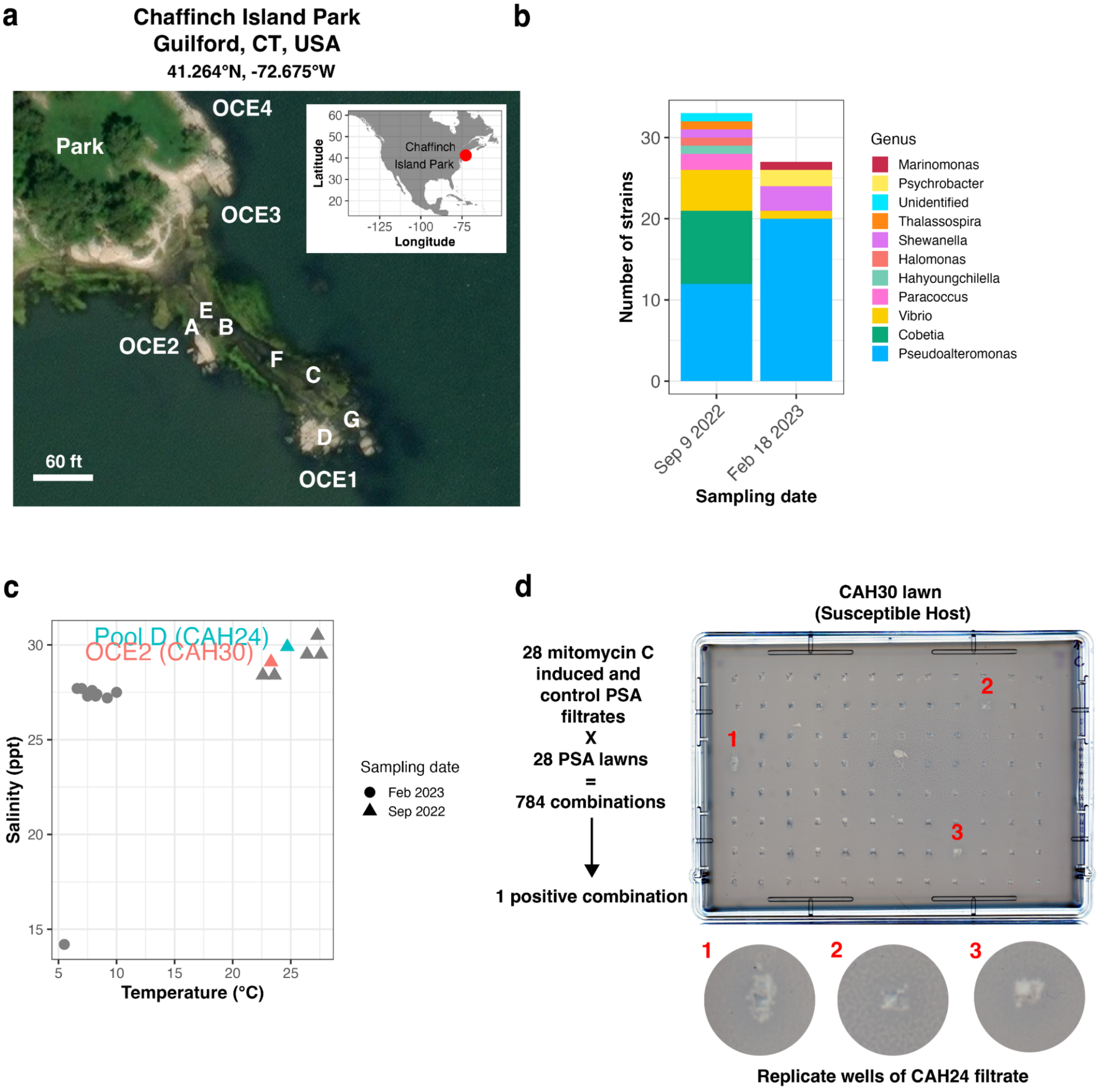

**Supp. Fig. 1. Isolation of marine strains from the northeastern USA included a *Pseudoalteromonas* isolate carrying an active prophage. (a)** Sampling locations at Chaffinch Island Park, Guilford, CT, USA. Single letters indicate tide pools, while “OCE” sites represent nearshore, upper surface waters of the Long Island Sound. Sites were sampled on September 9, 2022 and February 18, 2023 at approximately peak low tide, with sites A, B, C, D, and OCE1 sampled on both dates. Sites OCE2 and OCE3 were only sampled in September, and sites E, F, G, and OCE4 were only sampled in February. Aerial image was generated using Yale University ArcGIS Online. **(b)** Genus identification for isolates collected from sites in panel **a**, based on partial 16S sequencing of the V3-V4 region. Note that *Pseudoalteromonas* (PSA) was the most highly represented genus in this library. **(c)** Relationship between temperature (°C) and salinity (ppt) at our sampling sites on both dates, with labeled points representing sites that generated strains of interest in this manuscript. **(d)** Screen plate resulting from a mitomycin C induction assay, in which 28 PSA strains were incubated in the presence and absence of mitomycin C, and resulting filtrates spotted on lawns of all 28 strains. The marked wells indicate the location of the three triplicate wells of CAH24 filtrate on the CAH30 lawn.

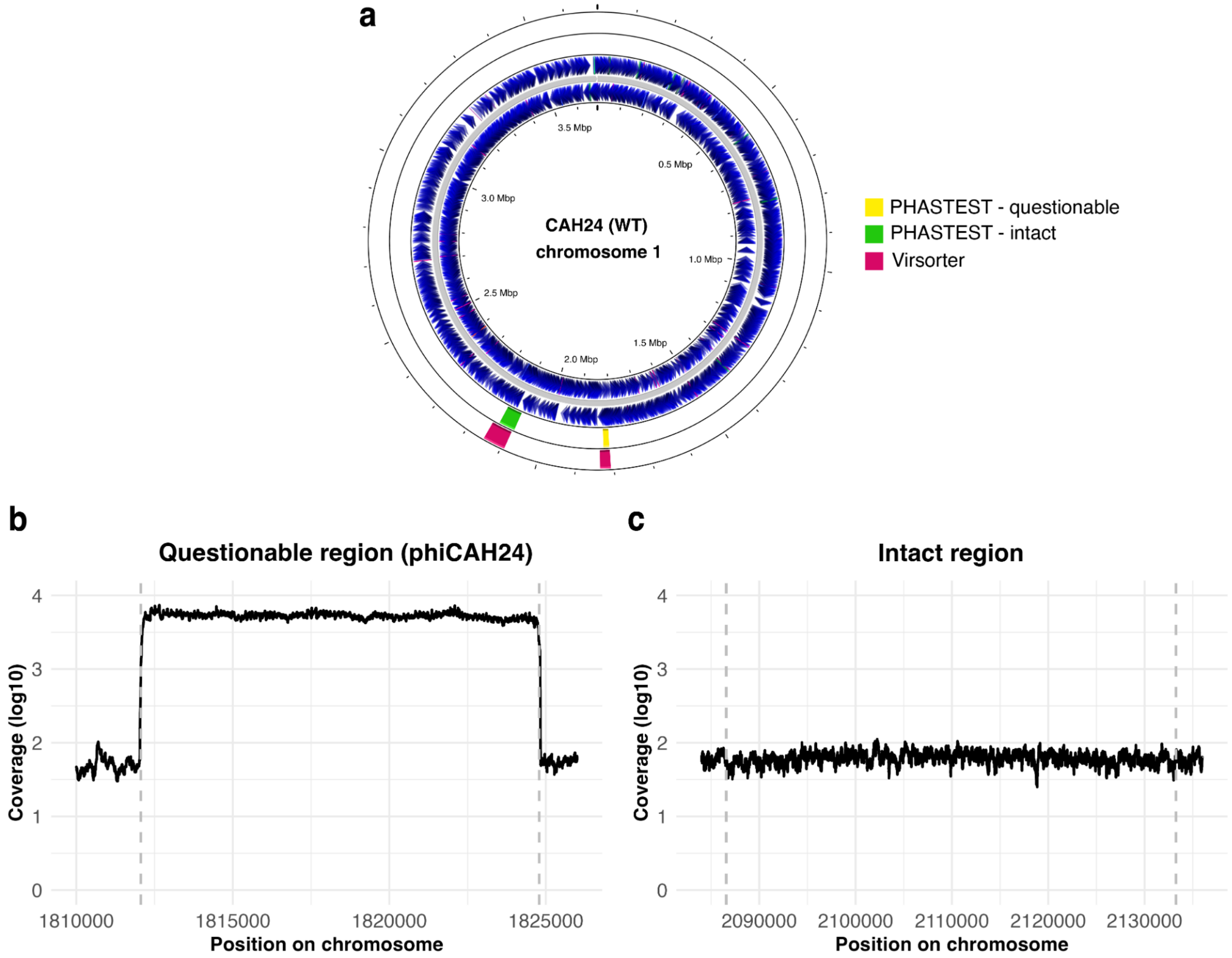

**Supp. Fig. 2. Read coverage of predicted prophage regions indicated that phiCAH24 is spontaneously active. (a)** Depiction of the CAH24 genome annotation of the major chromosome, visualized using Proksee. The genome is oriented to start at the predicted dnaA gene. Color blocks in the outermost ring indicate Virsorter predicted prophage regions, and the next inner ring indicates PHASTEST predicted prophage regions (with color representing completeness score). **(b)** Short read coverage of the PHASTEST “questionable” region (corresponding to phiCAH24) from sequencing of an overnight bacterial culture. Prophage genome boundaries are marked with a vertical dashed line, and were determined from subsequent sequencing of free viral particles. **(c)** Short read coverage of the PHASTEST “intact” region from sequencing of an overnight bacterial culture. Putative prophage boundaries are marked with a vertical dashed line, and are based on the PHASTEST prediction.

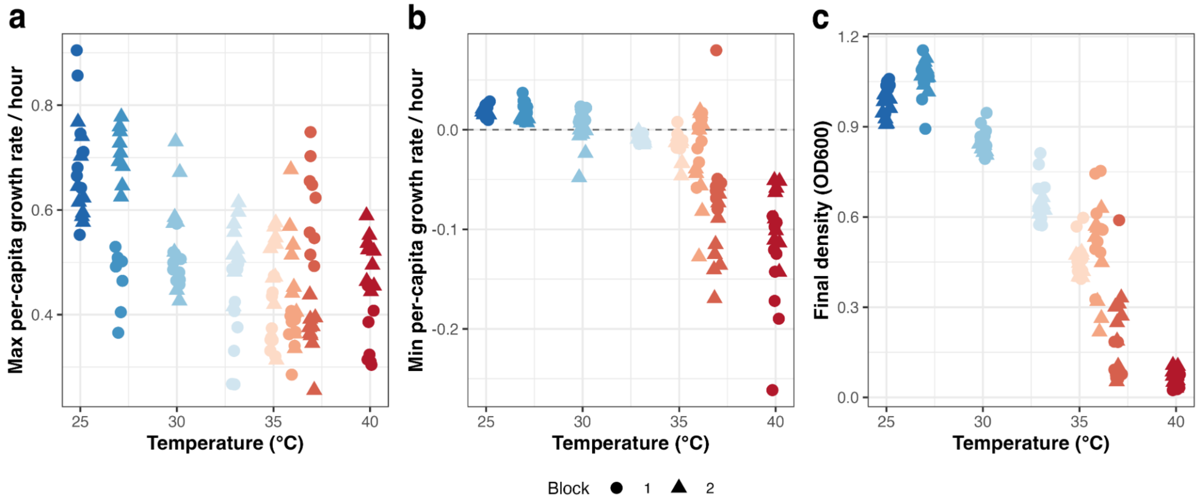

**Supp. Fig. 3. Growth rate and final density metrics for WT curves across temperatures.** Observed per-capita growth rates were calculated using the slopes of ln(OD600) with a 5 time point rolling window linear regression, and multiplying by a scaling factor of 4 to generate a per hour rate (given OD readings every 15 minutes). In each panel, individual points represent values generated from a single well at a given temperature within each block. **(a)** Maximum per-capita growth rate across each full growth curve. **(b)** Minimum per-capita growth rate after reaching a threshold OD of 0.07, to identify crashes. **(c)** Density (raw OD600, not transformed) at the end of each growth curve.

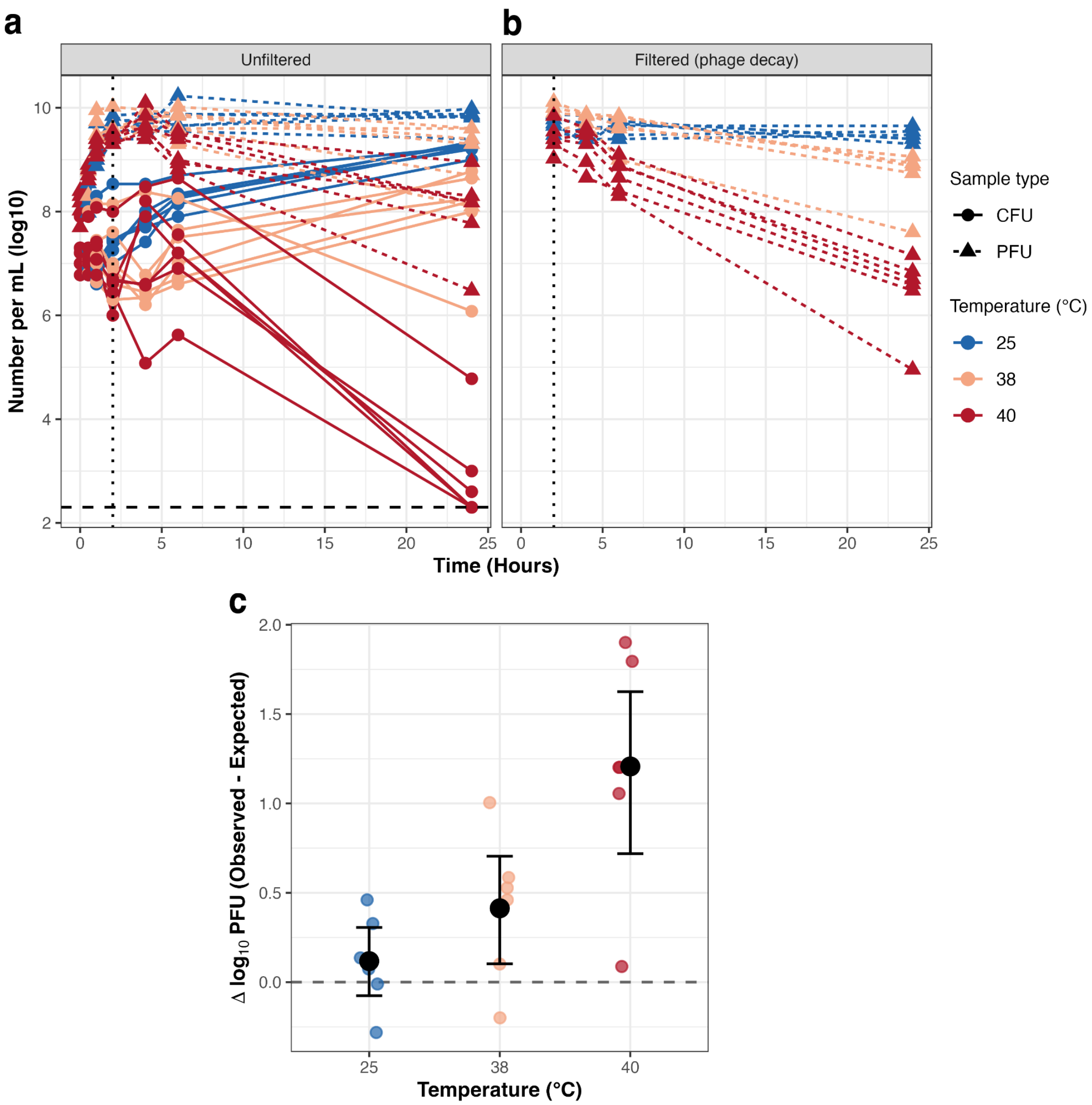

**Supp. Fig. 4. Wild type CAH24 bacteria and phage population dynamics and phage decay rates across temperature. (a)** Bacteria (CFU) and phage (PFU) densities over time across three temperatures. Lines connect samples derived from the same initial overnight culture, with each culture derived from a different WT bacterial colony. **(b)** Phage titers over time after filtration of cultures that had been incubated at a given temperature for two hours. Note that these samples were derived from the same initial cultures as in panel a, but were independent wells in the assay (i.e., sampling did not impact results in panel a). **(c)** Difference in final observed and expected PFUs given the estimated decay rates calculated from panel **b**. The large point represents the mean for each temperature, with error bars showing bootstrapped 95% confidence intervals.

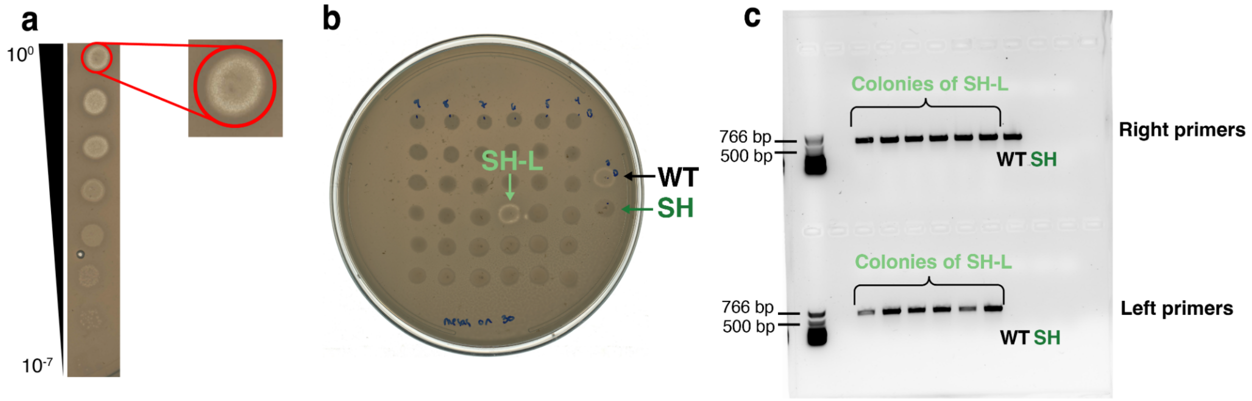

**Supp. Fig. 5. Generation and confirmation of prophage integration in the susceptible host background. (a)** A bacterial “mesa” growing within a high titer spot of phiCAH24 on a lawn of the susceptible host, CAH30. Note that this dilution series is the same as one depicted in Fig. 1a, and is used here simply as a representative example of a mesa. **(b)** Cultures generated from mesa-derived colonies were screened on a lawn of the susceptible host to identify putative phage-infected strains. Cultures were spotted in a 6x6 grid, with the WT (positive control) and SH (negative control) on the right side of the image. One mesa-derived culture had a visible ring of clearance (marked SH-L). **(c)** Agarose gel of PCR products using primers that span the host-phage junction on either side of the prophage, designed to amplify the novel lysogen genome. For each primer pair, one primer is within the prophage and the other is external on the host chromosome. Given that the WT strain is homologous to the SH on the right side of the prophage integration site (see Fig. 3a), that primer pair is expected to amplify both the WT and SH-L but not the uninfected SH strain. The left side primers are expected to uniquely amplify the novel lysogen SH-L.

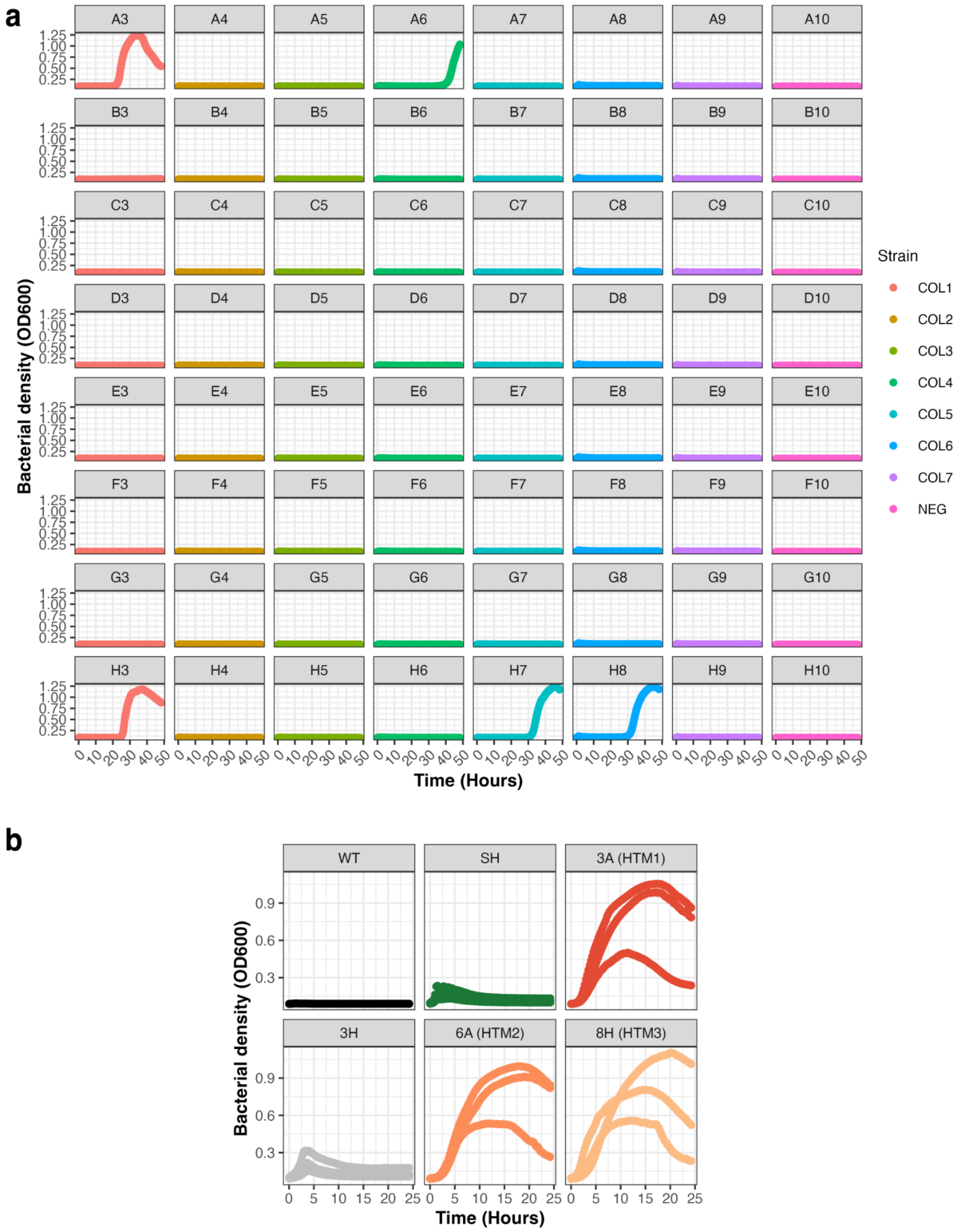

**Supp. Fig. 6. Growth curves of initial populations and restreaked isolates from the high temperature evolutionary rescue experiment. (a)** Bacterial density over time (OD600) of replicate populations of the wild type strain at a temperature just past this strain's upper thermal limit (38°C). Wells in each column (except the last, which is a negative control), were founded by an overnight culture from a different colony of the wild type CAH24 strain. **(b)** 38°C growth curves of isolates derived from turbid wells at the end of the initial experiment, with facet label specifying the initial well. The WT and SH strains were included for comparison. The isolates from wells 3A, 6A, and 8H demonstrated robust growth and were subsequently designated the “high temperature mutants” HTM1, HTM2, and HTM3, respectively.

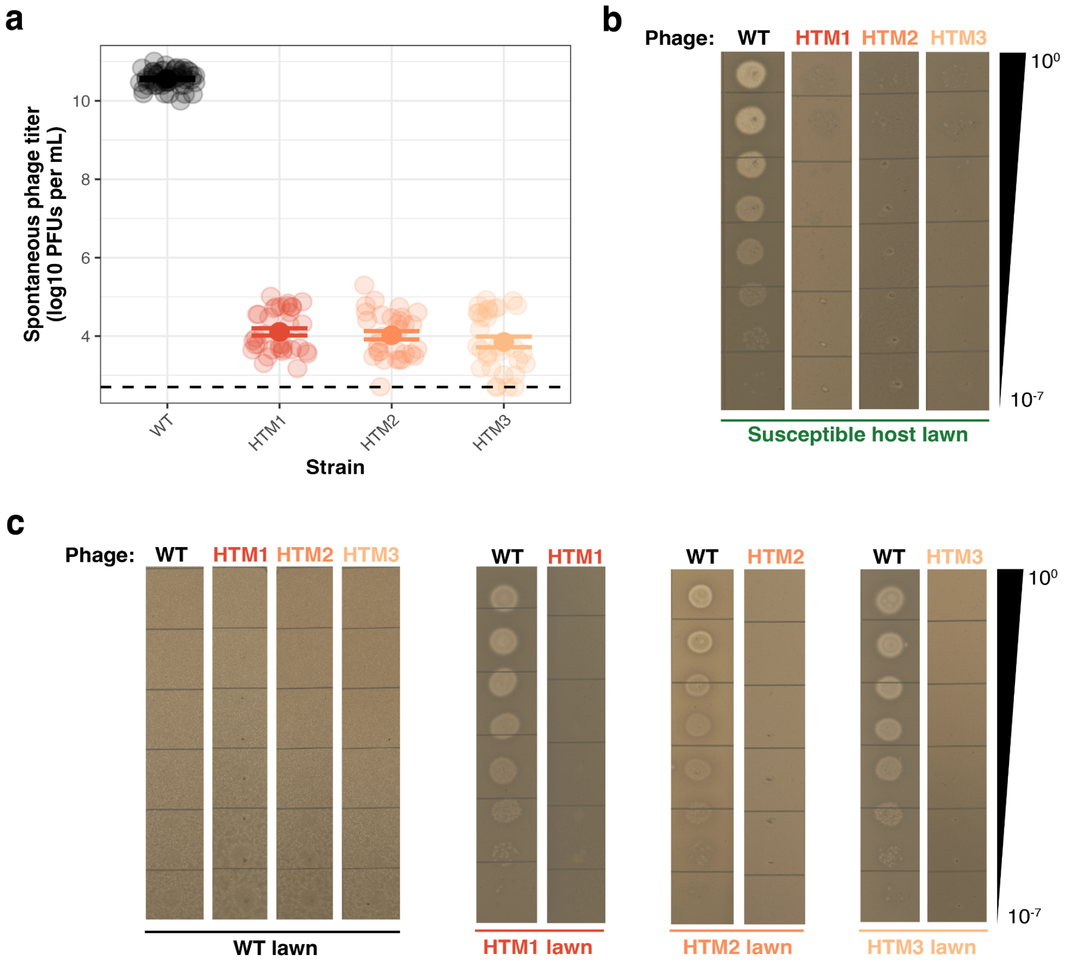

**Supp. Fig. 7. Spontaneous phage production and representative efficiency of plating images for the WT and HTMs. (a)** Spontaneous phage production titers of the WT and HTMs plated on the susceptible host. Each data point represents a unique combination of phage-producing colony and culture in the lawn, with each strain plated across at least six days. Data are summarized as mean ± SE. **(b)** Representative images of dilution series of WT and HTM-produced phage plated on the susceptible host. **(c)** Representative images of dilution series of WT and HTM-produced phage plated on WT and HTM lawns.

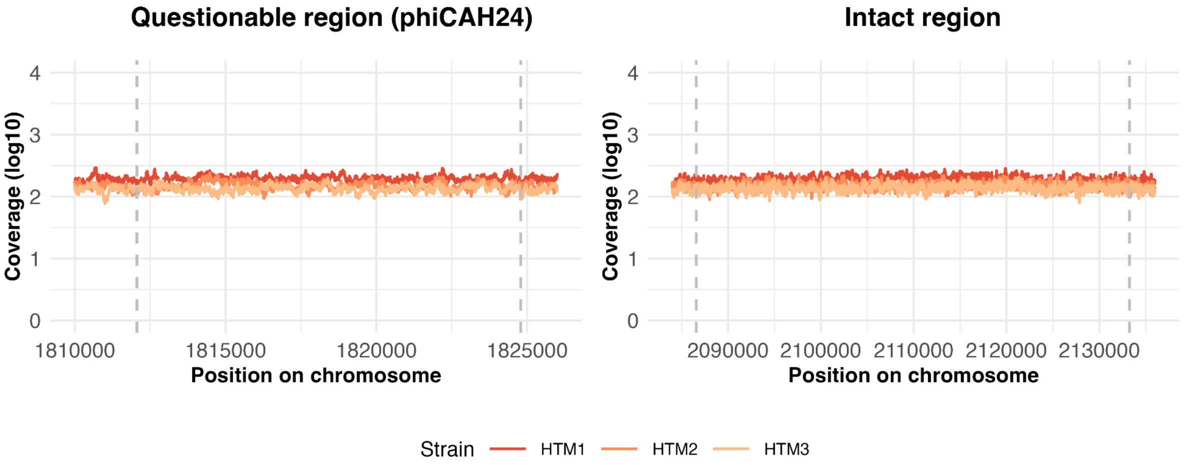

**Supp. Fig. 8. Read coverage in the predicted prophage regions from short read sequencing of the high temperature mutants.** Illumina short reads from overnight cultures of each HTM were aligned to the WT CAH24 reference genome, with predicted prophage boundaries indicated by the dashed lines.

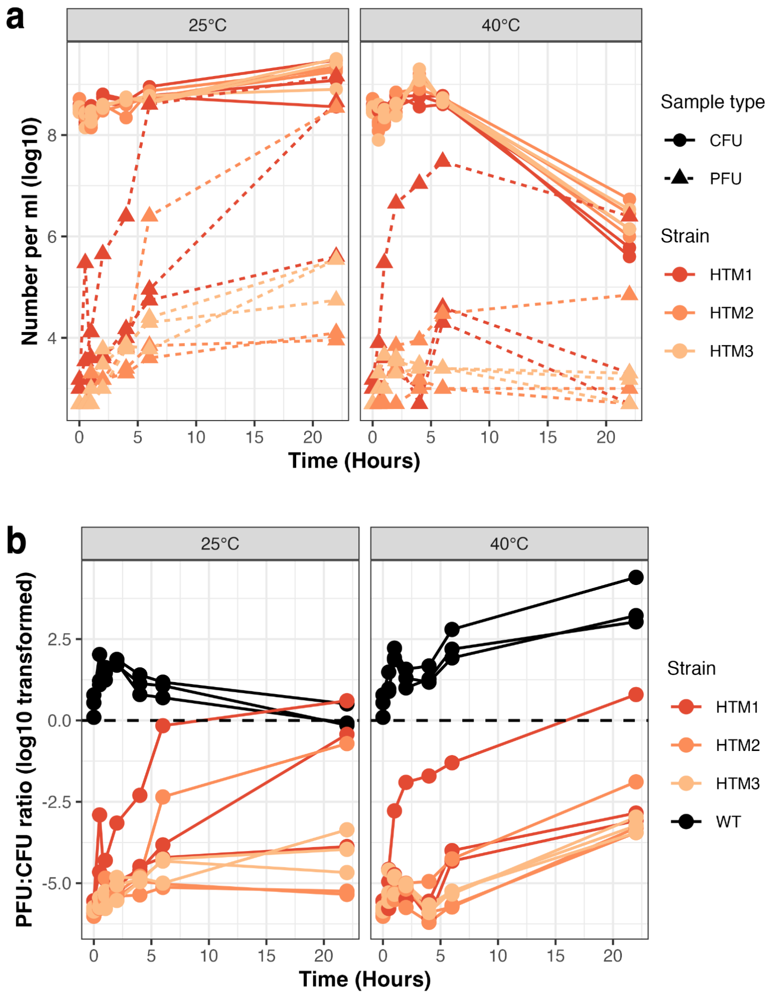

**Supp. Fig. 9. HTM bacteria and phage population dynamics at the two thermal extremes in this study. (a)** Bacteria (CFU) and phage (PFU) densities over time at 25°C and 40°C for each of the HTMs. Lines connect samples derived from the same initial overnight culture, with each culture derived from a different HTM colony. **(b)** PFU:CFU ratios over time calculated from the data in panel **a**, with the WT strain results shown for reference (same data shown in Fig. 2d).

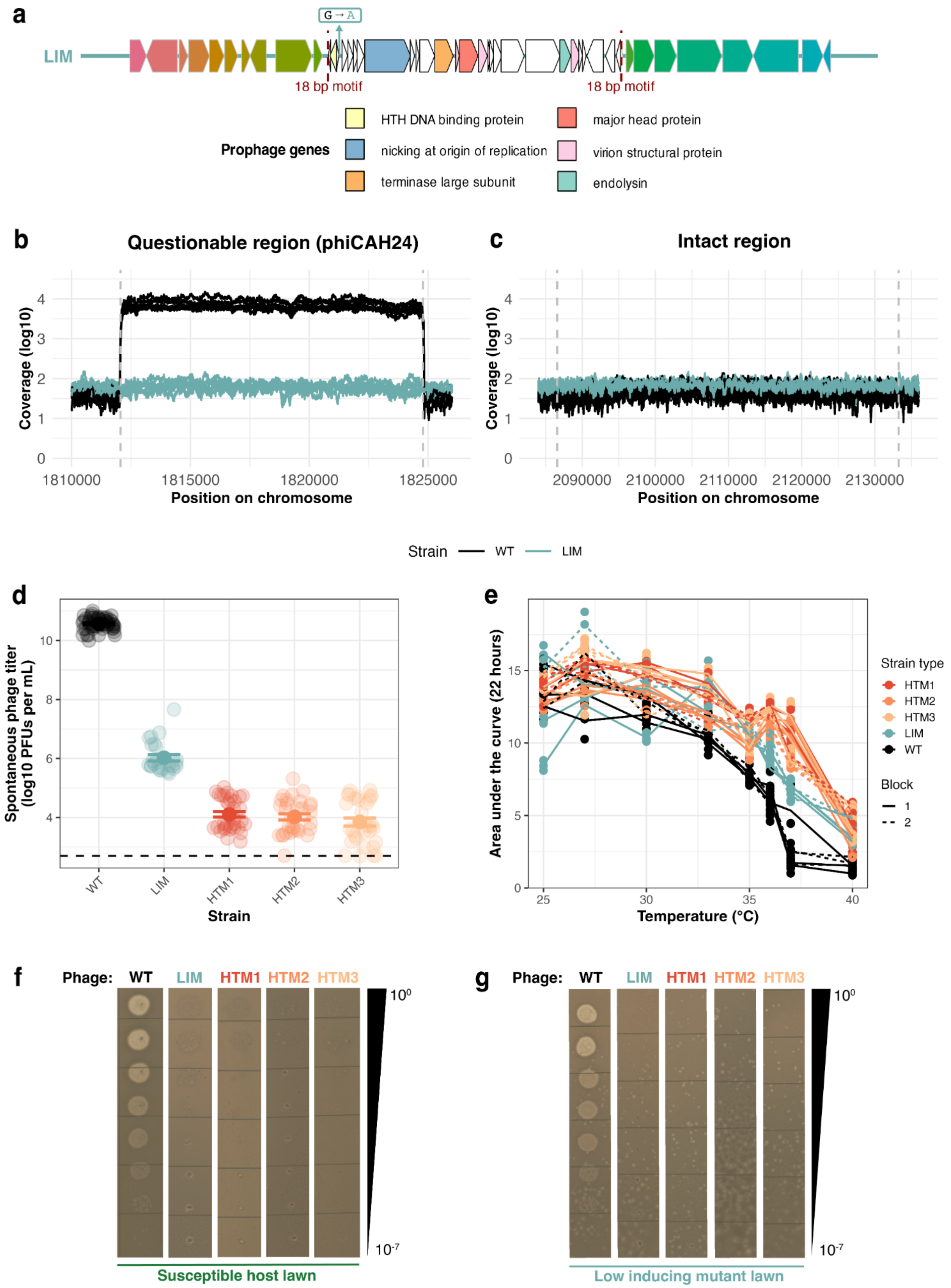
a
LIM
G→A
18 bp motif
18 bp motif
Prophage genes
HTH DNA binding protein
nicking at origin of replication
terminase large subunit
major head protein
virion structural protein
endolysin
b
Questionable region (phiCAH24)
Coverage (log10)
Position on chromosome
c
Intact region
Strain WT LIM
d
Spontaneous phage titer (log10 PFUs per mL)
Strain
WT LIM HTM1 HTM2 HTM3
e
Area under the curve (22 hours)
Temperature (°C)
Strain type
HTM1 HTM2 HTM3 LIM WT
Block 1 2
f
Phage: WT LIM HTM1 HTM2 HTM3
Susceptible host lawn
g
Phage: WT LIM HTM1 HTM2 HTM3
Low inducing mutant lawn

**Supp. Fig. 10. Opportunistic discovery of low inducing mutant (LIM) colonies demonstrates the consequences of additional mutations in the HTMs. (a)** Location of the single shared SNP found in all LIMs relative to the WT genome, found in an intergenic region within the prophage. **(b)** Short read coverage of the phiCAH24 region in overnight cultures of the WT and LIMs. Six independent colonies of each strain were used to generate these reads. **(c)** Short read coverage in the second predicted prophage region. **(d)** Phage titers spontaneously produced by overnight cultures of the WT, LIMs, and HTMs, plated on the SH. WT and HTM data are the same as shown in Supp. Fig. 7 and are shown for comparison. Each point represents an independent combination of phage-generating culture and SH colony in the lawn, with each strain plated across at least five days. Data are summarized as mean ± SE. **(e)** Thermal reaction norm using the area under the curve for each strain across temperature. Points represent the individual values for a given growth curve well, with lines connecting the means for the same colony at each temperature. **(f)** Representative images of dilution series of WT, LIM, and HTM-produced phage plated on a lawn of the SH. **(g)** Representative images of dilution series of WT, LIM, and HTM-produced phage plated on a LIM lawn.

### Supplementary Tables

**Supp. Table 1. Assembly statistics for the WT and SH genomes.** Values were determined by using Bowtie 2 to map Illumina short reads to the output of the Unicycler hybrid assemblies, with assembly statistics quantified using samtools.

| Strain | Contig | Contig Size | Coverage | Mean Depth | Mean Base Quality | Mean Map Quality |
|---|---|---|---|---|---|---|
| **CAH24 (WT)** | 1 | 3.69 Mb | 100 | 81.7 | 32.7 | 41 |
| | 2 | 788 kb | 100 | 74.2 | 32.7 | 41.4 |
| | 3 | 202 kb | 99.9995 | 88.3 | 32.7 | 41.4 |
| **CAH30 (SH)** | 1 | 3.69 Mb | 100 | 108.5 | 32.6 | 40.9 |
| | 2 | 797 kb | 100 | 108.6 | 32.6 | 41.4 |

**Supp. Table 2. Primers used in this study.**

| Primer name | Sequence | Amplicon size (bp) | Goal |
|---|---|---|---|

| | | | |
|---|---|---|---|
| **331F** | TCCTACGGGAGGCAGCAGT | 466 | Identifying isolates using the V3-V4 region of 16S |
| **797R** | GGACTACCAGGGTATCTAATCCTGTT | | |
| **Right_F** | TATCGTCAAAACCAAGACCAG | 566 | Amplifying the right region of the prophage boundary in the novel lysogen. Expected to also amplify the WT but not the SH. |
| **Right_R** | GCCCGAAACTCATAAAACCA | | |
| **Left_F** | TGGTATAGCAAAACAAGCCC | 648 | Amplifying the left region of the prophage boundary in the novel lysogen. Not expected to amplify the WT or SH. |
| **Left_R** | AAAGATTGCTCGCTGTGAAT | | |

138
